# SARS-CoV-2 Nsp1 N-terminal and linker regions as a platform for host translational shutoff

**DOI:** 10.1101/2022.02.10.479924

**Authors:** Andrea Graziadei, Fabian Schildhauer, Christian Spahn, Matthew Kraushar, Juri Rappsilber

## Abstract

In the early stages of SARS-CoV-2 infection, non-structural protein 1 (Nsp1) inhibits the innate immune response by inserting its C-terminal helices into the mRNA entry channel of the ribosome and promoting mRNA degradation. Nevertheless, the mechanism by which Nsp1 achieves host translational shutoff while allowing for viral protein synthesis remains elusive. We set out to characterize the interactome of full-length Nsp1 and its topology by crosslinking mass spectrometry in order to investigate the role of the N-terminal domain and linker regions in host translational shutoff. We find that these regions are in contact with 40S proteins lining the mRNA entry channel and detect a novel interaction with the G subunit of the eIF3 complex. The crosslink-derived distance restraints allowed us to derive an integrative model of full-length Nsp1 on the 40S subunit, reporting on the dynamic interface between Nsp1, the ribosome and the eIF3 complex. The significance of the Nsp1-eIF3G interaction is supported by further evidence that Nsp1 predominantly binds to 40-43S complexes. Our results point towards a mechanism by which Nsp1 is preferentially recruited to canonical initiation complexes, leading to subsequent mRNA degradation.

## Introduction

Coronaviruses (CoVs) are positive-sense single-stranded mRNA viruses capable of infecting a large variety of vertebrate species, causing mild to severe respiratory diseases, including severe acute respiratory syndrome (SARS). SARS-CoV-2 is the causative agent of the highly pathogenic respiratory disease COVID-19. It belongs to the beta genus of CoVs that includes SARS-CoV and the Middle East respiratory syndrome (MERS) CoV.

Beta-CoVs extensively remodel cell morphology (Cortese *et al*, 2020) and gene expression upon infection in a time-dependent manner (Finkel *et al*, 2021; Kamel *et al*, 2021) by a combination of viral proteins, transcripts and selective activation of host cell pathways. The hijacking of protein synthesis is a key phenotype of CoV infection, leading to ribosomes selectively translating viral factors and some host proteins. Shortly after internalization, SARS-CoV employs its non-structural protein 1 (Nsp1) to bind the small ribosomal subunit and inhibit canonical mRNA translation, resulting in cellular mRNA degradation (Lokugamage *et al*, 2012; Kamitani *et al*, 2009). This occurs via an unknown mechanism involving downstream endonucleolytic cleavage of host mRNAs (Tanaka *et al*, 2012; Kamitani *et al*, 2009). The resulting suppression of host protein synthesis and immune response includes the translation-dependent type I interferon pathway (Narayanan *et al*, 2008), a phenotype known as host translational shutoff.

CoV-2 Nsp1, which shares 84% sequence identity with CoV Nsp1 (Yuan *et al*, 2021), has been identified as a binder of the 40S ribosome in both biochemical and structural work (Schubert *et al*, 2020; Thoms *et al*, 2020). The protein has an N-terminal domain (NTD), a linker region and a small C-terminal domain (CTD) made up of two short alpha helices. Both CoV and CoV-2 Nsp1 can induce cleavage of capped and IRES-containing mRNAs *in vitro (Huang et al, 2011; Lokugamage et al, 2012; Mendez et al, 2021)*.

Recent cryo-electron microscopy (cryo-EM) structures have illustrated the mechanism by which the Nsp1 CTD renders the 40S and 80S ribosomes, and several 43S initiation complexes, translationally incompetent (Schubert *et al*, 2020; Thoms *et al*, 2020). The helices making up the C-terminal domain of Nsp1 plug the mRNA entry channel on the 40S subunit, preventing translation. However, only the C-terminal domain and a short portion of the linker were resolved in the structures, presumably due to flexibility of the rest of the protein. Biochemical work has since then highlighted the role of the N-terminal domain and the linker in viral evasion of translational shutoff (Shi *et al*, 2020; Tanaka *et al*, 2012; Tidu *et al*, 2020). However, the mechanism by which this occurs remains unknown.

Here, we characterize the conformation of full-length Nsp1 on the 40S and 43S complexes by crosslinking mass spectrometry, and probe its network of protein-protein interactions. Crosslinking MS shows that the previously unresolved Nsp1 NTD, while highly flexible, occupies a well-defined volume on the 40S subunit, and that its linker contributes to interactions with the 40S, contextualizing previously described clinical variants. Using the crosslinking MS data, we build an integrative model combining current structural knowledge with spatial restraints for Nsp1. We then characterize the Nsp1-bound proteome by affinity purification mass spectrometry (AP-MS) and ultracentrifugation. Taken together, our results show that Nsp1 preferentially binds to 40S subunits and 43S preinitiation complexes, and that both the linker and NTD regions make extensive interactions with the 40S subunit and eukaryotic initiation factors (eIFs). The data point to a role of the eIF3 subunit eIF3G, which can be unambiguously assigned to be in close proximity to Nsp1 near the mRNA entry site based on crosslinks, clarifying the identity of the protein occupying low-resolution density in previous structures (Thoms *et al*, 2020). We hypothesize that these dynamic interactions by the Nsp1 linker and NTD to initiation factors contribute to the preferential targeting of canonical initiation complexes, and that translation on the viral 5’ UTR may proceed in an eIF3G-independent manner.

### The flexible SARS-CoV-2 Nsp1 NTD and linker on the 40S subunit

In order to understand the topology of the protein-protein interaction network involved in Nsp1-mediated translational repression, we performed a crosslinking-MS analysis of HEK293T cells overexpressing FLAG-tagged Nsp1. After cell lysate was crosslinked with disuccinimidyl sulfoxide (DSSO), the Nsp1-bound proteome was enriched by affinity purification, digested and analyzed by crosslinking-MS. We identified 1,269 crosslinks involving the same protein (self-links) and 869 heteromeric crosslinks at a 2% residue-pair false discovery rate (FDR), resulting in the identification of 515 protein-protein interactions (PPIs) at a 6% FDR. Identified crosslinks covered the ribosome, signal recognition particle, chaperonin-containing T-complex, as well as eIF1, eIF3, eIF2, and eIF4F complexes. As crosslinks directly report on the proximity of residues, we could map the binding site of Nsp1 on ribosomes in solution (Fig. 1). This was consistent with the arrangement proposed by cryo-EM structures, though several additional interactions were detected involving the regions of Nsp1 not previously resolved. We detected 42 crosslinks between Nsp1 NTD and linker to 40S ribosomal proteins. These include a previously reported interaction with RS2 (uS5) and RS3 (uS3) (Slavin *et al*, 2021; Thoms *et al*, 2020; Schubert *et al*, 2020), and novel contacts with RS9 (uS4), RS10 (eS10), RS12 (eS12), RS17 (eS17), and RS27A (eS31), which line the cavity near the mRNA entry site. While no crosslinks between the Nsp1 CTD and the ribosome were observed, this is likely due to the fact the CTD has a single lysine residue that lies away from RS2 and RS3 (Fig. S3C), though it is possible that Nsp1 binds a subpopulation of the ribosomes in the dataset without plugging the mRNA entry site with its CTD (see discussion).

**Fig. 1.**
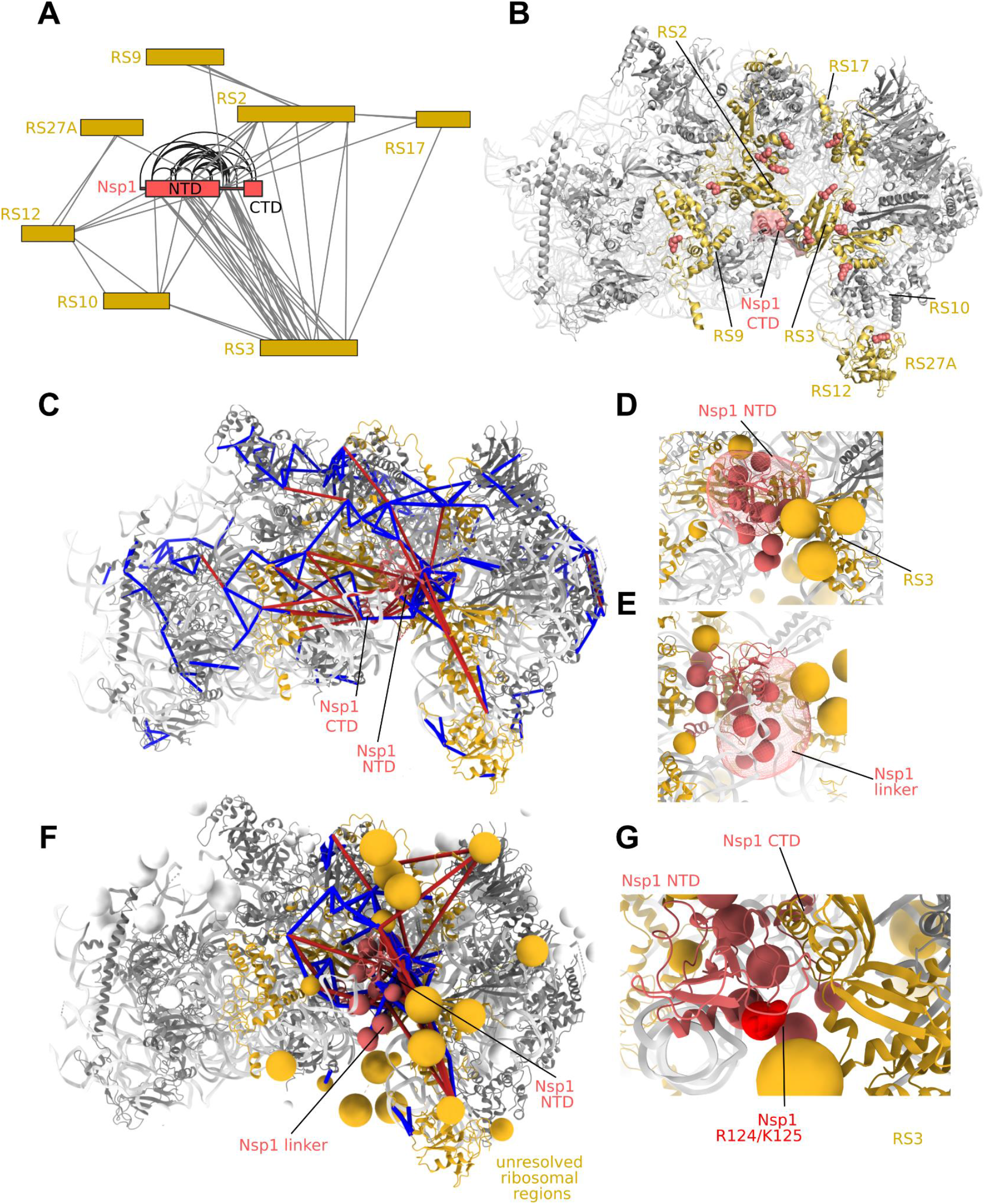
Protein-protein interactions of the Nsp1 NTD. **A)** Interactors of Nsp1 by crosslinking-MS. The Nsp1 NTD and linker regions are observed in a large number of interactions with ribosomal S3 (uS3). Crosslinks within Nsp1 are also displayed. **B)** 40S subunit (pdb 6zlw) proteins interacting with Nsp1 highlighted in gold. Residues showing crosslinks to Nsp1 shown as spheres. **C)** Integrative modeling of Nsp1 NTD onto the 40S ribosome. The solution represents the centroid of the most populated cluster of models, with the Nsp1 NTD interacting with RS3. Crosslinks are mapped onto the complex. Satisfied crosslinks (<30Å) in blue, violated crosslinks are in red. The crosslinks show secondary populations where the Nsp1 NTD samples the other side of the cavity facing the mRNA entry site. **D)** Localization probability density of the Nsp1 NTD in the main cluster of models. Beads comprising the coarse grained regions not present in the structure (Nsp1 linker, flexible ribosomal protein regions) are shown. **E)** Localization probability density for the Nsp1 linker region in the main cluster. The linker largely localizes at the bottom of RS3. **F)** Integrative modeling result with all coarse grained regions displayed. All Nsp1 crosslinks shown, including those to coarse grained regions. **G)** In the main cluster of models, K125 is observed in close proximity to RS3.

We used the restraints derived from crosslinking-MS data to derive a model of the interactions of full-length Nsp1 on 40S ribosomal subunits using the integrative modeling platform (IMP) (Russel *et al*, 2012) (Fig. S1). A coarse-grained representation of the Nsp1-bound 40S initiation complex, including all regions not resolved in structures, was used in the derivation of models representing conformations consistent with crosslinking and physical restraints. The Nsp1 NTD, linker and 40S unresolved regions were kept fully flexible with respect to the Nsp1 CTD-ribosome body.

In the models, Nsp1 NTD occupies a well-defined volume in the cavity facing the mRNA entry site, interacting with S3 (Fig. 1C, 1D, S3, S4). Nevertheless, multiple positions of the NTD are consistent with crosslinking restraints (Fig. S3), indicating the dynamic nature of the Nsp1 NTD on the 40S complex and explaining why it had not been resolved in previous structures. In particular, several crosslinks identify a secondary extended conformation in which the Nsp1 NTD lies in proximity to RS10 and RS12 (Fig. 1C).

The Nsp1 linker region has multiple crosslinks to RS2, RS3, RS10 and RS12. The localization probability density of the linker in our final model indicates that the linker is largely restricted to the right side of the cavity, localizing below the plane of the Nsp1 NTD. Indeed, the recent observations that the NTD and linker regions are required for translational suppression (Shi *et al*, 2020) are consistent with our findings that the linker is not a passive bystander of Nsp1-ribosome recognition, but rather makes extensive and dynamic contacts with the cavity facing the mRNA entry channel.

### The interactome of Nsp1 indicates a targeting of translation initiation

We further characterized the nature of the Nsp1-bound proteome by affinity purification mass spectrometry (AP-MS) experiments in HEK293T cells. Quantitative AP-MS analysis identified 271 high-confidence co-purifying proteins (Fig. 2, supplementary table S3), observing good correlation between experimental replicas (R=0.9-0.95, Figure S6). As expected, ribosomal proteins made up a majority of co-purifying proteins.

**Fig. 2.**
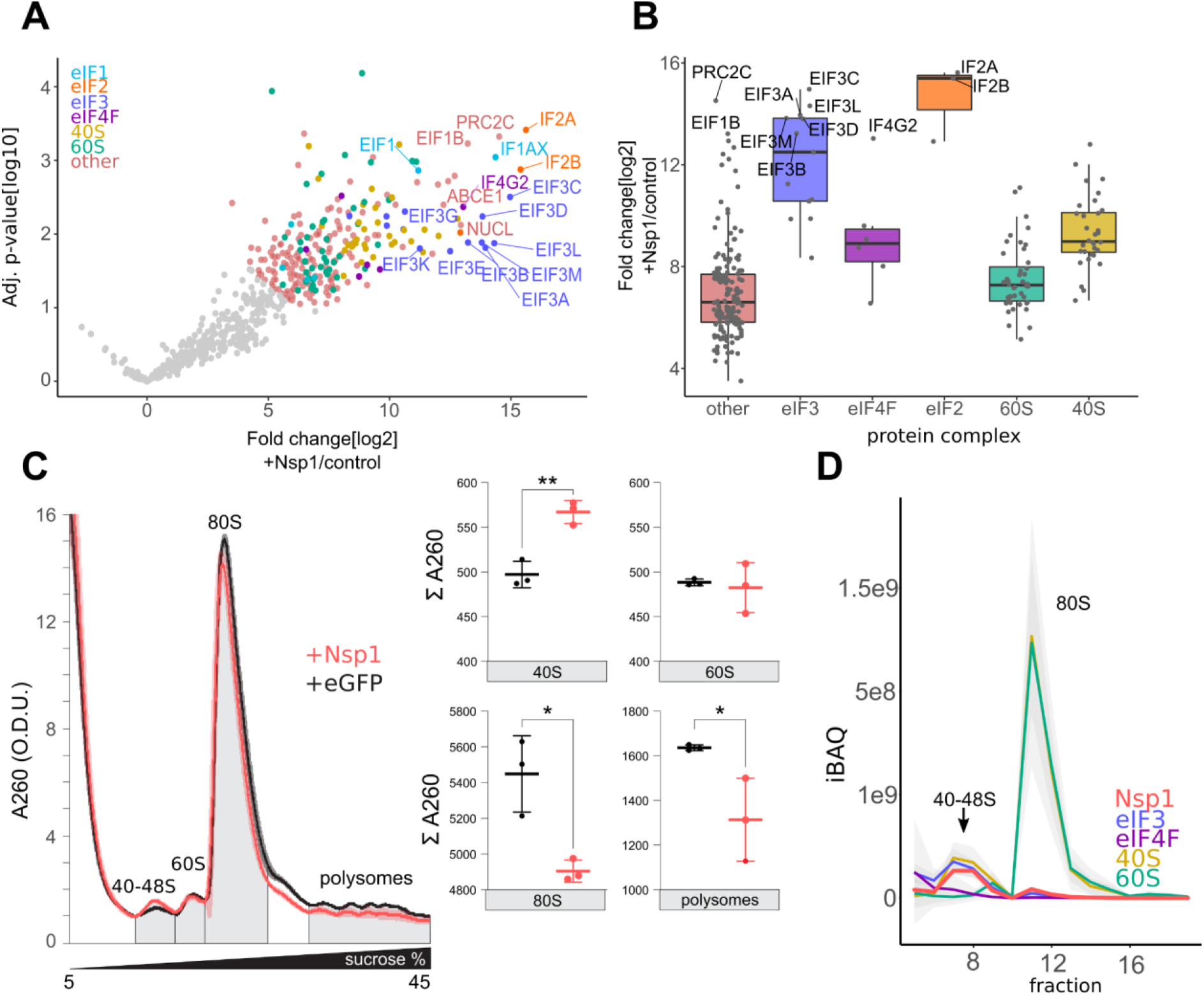
Nsp1 preferentially targets initiation complexes. **A)** Volcano plot showing the log2 fold change and the significance (adjusted p value) of enriched proteins in affinity purification pull-downs of 3xFLAG-tagged Nsp1 expressed in HEK293T cells. Significantly enriched (FDR<0.05) proteins are highlighted and coloured by protein complex. Proteins are labelled according to their Uniprot Entry name. Subunits of the eIF3 complex, as well as eIF2α (IF2A) and eIF2β (IF2B) are highly enriched. **B)** Fold enrichment of significantly enriched protein complexes in AP-MS data. The only subunit of the eIF4F complex to be highly enriched is the non-canonical eIF4G2 (Dap5). **C)** Left: Analytic density gradient fractionation of A260-normalized HEK293T lysates overexpressing Nsp1 or eGFP control. The absorbance traces measure the relative abundance of 40-48S complexes, 60S subunits, 80S ribosomes, and polysomes. A260 curves plotted as means of 3 replicates, with the shaded area covering 1 standard deviation. Right: Comparisons of area under the curve for Nsp1 and eGFP control. Student’s t-test: *, p <0.05, ** p<0.01. **D)** Proteomic co-fractionation analysis of analytic density gradients of cells expressing Nsp1. Nsp1 largely migrates in the 40-48S region, rather than in full ribosomes. It co-migrates with eIF3, eIF4F and 40S subunit proteins. The average iBAQ for each complex is shown, with shaded area covering 1 standard deviation of the abundances of each component.

Gene Ontology (GO) enrichment analysis showed translation initiation, overall translation and several terms associated with ribosome biogenesis among the most highly enriched terms (Fig. S7). These findings are consistent with literature showing Nsp1-mediated shutdown of host mRNA-translating ribosomes in the early stages of infection (Kamel *et al*, 2021; V’kovski *et al*, 2021; Finkel *et al*, 2021).

Translation initiation factors were among the most highly enriched proteins in the dataset, surpassing even ribosomal subunits that were identified by structural models as direct Nsp1 binders. In addition to factors identified in cryo-EM studies (LYAR, PAIRB, TSR1) (Thoms *et al*, 2020; Schubert *et al*, 2020), we also detected subunits of the exosome complex and the exon junction complex, possibly indicating the nature of downstream mRNA pathways acting on mRNA-stalled ribosomes. This is also consistent with reports of nuclear activity of Nsp1 (Zhang *et al*, 2021).

The top hit by fold enrichment in the AP-MS screen is eIF2α (EIF2S1/IF2A), a key player in the integrated stress response pathway (Clemens, 2001), whose phosphorylation by stress-sensing kinase leads to suppression of translation initiation. Phosphorylation of eIF2α is known to be inhibited by beta-CoV factors such as the MERS 4a protein (Rabouw *et al*, 2020, 2016). Interestingly, the interferon-inducible activator of the EIF2AK2 stress-sensing kinase, PRKRA, is also detected as significantly enriched in the AP-MS screen, along with the relevant type I interferon pathway protein DDX60, although these proteins may also co-purify with Nsp1 and Nsp1-associated ribosomes via interactions with the mRNA.

In addition to those initiation factors found in previously described cryo-EM structures (eIF3, eIF2, eIF1, eIF1AX) (Thoms *et al*, 2020; Schubert *et al*, 2020), other canonical components of the 43S initiation complex were detected, such as eIF1B and the helicase DHX29. The subunits of the eIF4F complex, eIF4B and 4IF4H were also found to be enriched. These proteins are involved in unwinding structures in the 5’ UTR region and mediating the attachment of the 43S complex to the mRNA, enabling scanning upon ATP hydrolysis (Jackson *et al*, 2010; Querido *et al*, 2020; Hinnebusch, 2014). Intriguingly, the polyA binding protein (PABP), an integral part of the eIF4F/eIF4B-mRNA complex that is loaded onto the 43S initiation complex to form the closed loop 48S complex, is not among the enriched proteins. Similarly, core components of eIF4F are less enriched than other eIFs, while the non-canonical initiation factor eIF4G2 (Dap5), is highly enriched (Fig. 2B). Thus, Nsp1 seems to be stabilizing late-stage 43S intermediates, as well as to be co-purifying with eIF4F/eIF4B. This interaction may occur on the ribosome itself, or via a previously unreported interaction outside the ribosome. The eIF2 guanosine exchange complex eIF2B was also detected among the enriched translation initiation factors.

The high abundance of translation initiation factors in our data led us to hypothesize that Nsp1 binding to ribosomes is not purely mediated by the interactions between the C-terminal helices and the mRNA entry site, which are shared between late-stage initiation complexes and elongating 80S ribosomes. We therefore analyzed the distribution of ribosomal complexes in cells overexpressing Nsp1 by sucrose gradient centrifugation. Here, we could observe a difference between the subunit distributions of cells overexpressing EGFP versus Nsp1 (Fig.2), which reproducibly showed an accumulation amount of free 40S subunit and 43-48S complexes for Nsp1.

Proteomic analysis of the sucrose gradient fractions confirmed that Nsp1 preferentially binds pre-initiation intermediates. This co-fractionation analysis indicated that Nsp1 is mainly found bound to 40S subunits and 43S ribosomal initiation intermediates (Fig. 2D). Interestingly, only a small fraction of Nsp1 fractionated with 80S ribosomes. Instead, Nsp1 showed coelution behavior with subunits from the eIF3 complex (Fig. S8) to which we had found extensive crosslinks. Moreover, the decrease of 80S ribosomes in the Nsp1-overexpressing cells did not lead to a corresponding 60S increase (Fig. 2C), consistent with preferential blockage of specific initiation intermediates over the bulk actively translating ribosome and polysome population.

### Structural proteomics of SARS-CoV-2 Nsp1

Consistent with the enrichment of eIF3 components in Nsp1 AP-MS, we observed extensive crosslinking between the Nsp1 linker and N-terminal domain regions and the G subunit of the eIF3 complex. With 18 detected crosslinks, eIF3G was detected as the primary interactor for Nsp1 in the sample. Using crosslinks, we could identify that the unidentified RNA-recognition motif (RRM) domain found near the mRNA entry site in cryo-EM structures (Thoms *et al*, 2020) is the RRM domain of eIF3G. This is also consistent with the identification of this domain in the structure of the 48S complex (Querido *et al*, 2020). The RRM domain is connected by a linker to the N-terminal region of eIF3G, which we could model positioned across the surface of the RNA-entry site (Figure 3C).

**Fig. 3.**
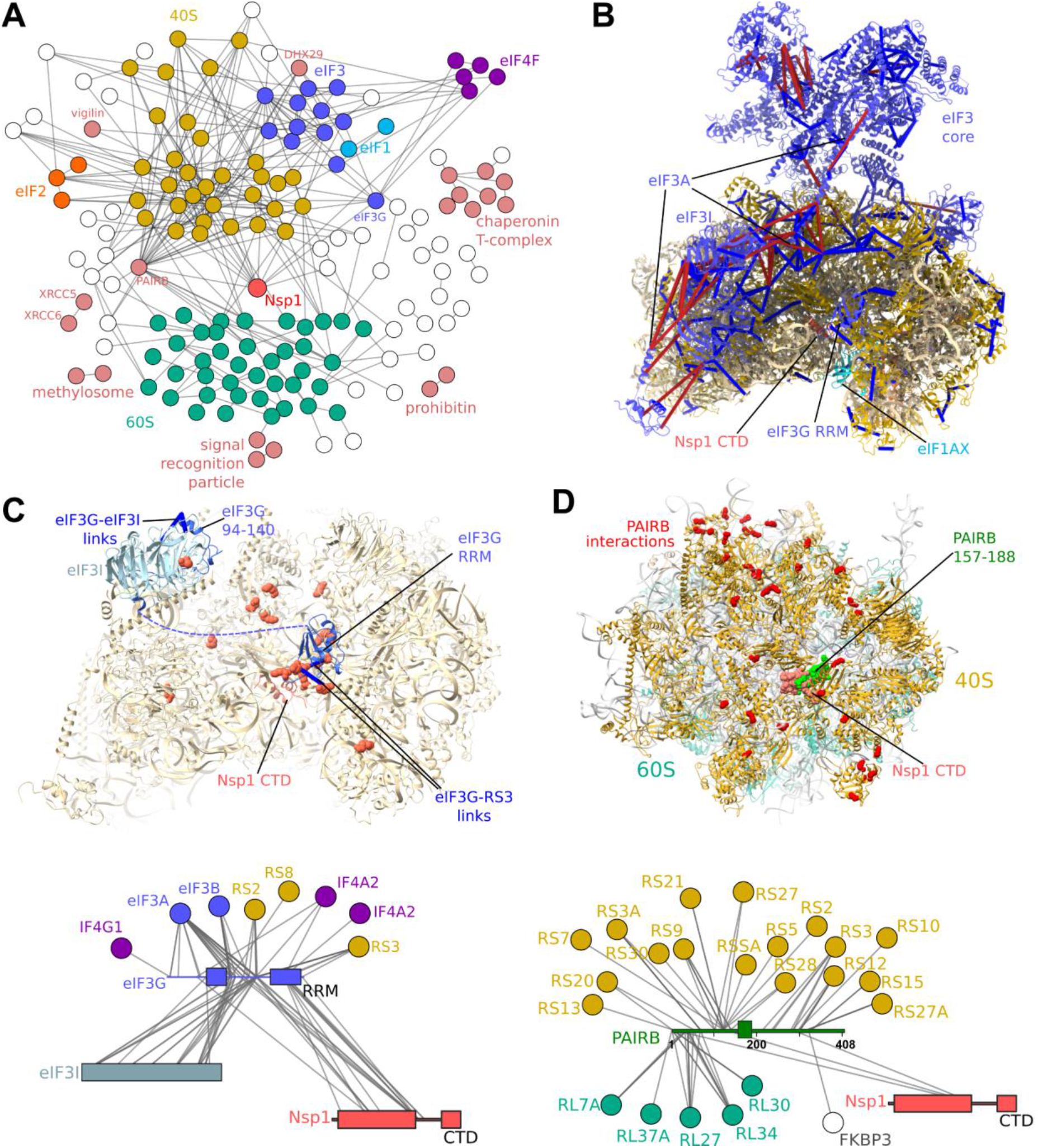
Protein-protein interaction map of the Nsp1-bound proteome. **A)** Crosslinking-MS detects 515 protein-protein interactions (PPIs) at a 6% PPI-level FDR. Each edge represents one or more crosslinks. Nsp1 primarily interacts with the 40S and eIF3 complexes. Additional interactions between the ribosome and regulatory or initiation factors are detected. **B)** Crosslinks mapped onto the structure of the Nsp1-bound 43S initiation complex (pdb 6zp4) (Thoms *et al*, 2020). Satisfied crosslinks (<30Å) in blue, violated crosslinks are in red. The crosslinks are largely consistent with the structure, with violations highlighting the mobility of the head region of eIF3, and of the eIF3I subunit. Crosslinks indicate a possible alternative register for the long helix of eIF3A, which was poorly resolved in the structure. **C)** Crosslink-guided assignment of the unidentified RRM domain in the complex, and modeling of the eIF3I-binding region of eIF3G by homology and crosslinking-MS data. 40S and eIF3G-RRM residues crosslinked to Nsp1 NTD and linker regions are shown as red spheres. **D)** PAIRB binds Nsp1-stalled ribosomes and 40S subunits View of the face of the 40S subunit with residues crosslinked to PAIRB shown as red spheres. Crosslinks show the large, disordered protein wraps around the 40S subunit, with its plug region displaced by the Nsp1 helices. Crosslinks mapped onto PAIRB-bound 80S ribosome (pdb 6z6n) (Wells *et al*, 2020). Crosslinks also show interactions between Nsp1 NTD and PAIRB.

While most crosslinks are between Nsp1 and eIF3G linker regions, structural visualization of the crosslinks on the eIF3G RRM domain showed that residues crosslinked to Nsp1 are positioned on the solvent-exposed face of the domain, consistent with their interaction occurring within the 43S complex. These multiple interactions are in line with the co-fractionation behavior of the two proteins (Fig. 2D, Fig. S8). Taken together, AP-MS and crosslinking-MS provide evidence for a direct or RNA-mediated interaction between Nsp1 and eIF3G in the context of late 43S initiation complexes. This is further bolstered by the detection of an extensive network of crosslinks involving eIF3G and Nsp1 K125, a residue that is required for translational repression and mRNA depletion (Mendez *et al*, 2021).

As indicated by previous structural work and by our proteomic analysis of ribosome preparations, Nsp1 can bind multiple ribosomal states all the way up to late-stage 43S initiation complexes. The eIF3G RRM covers up RS3, thus precluding some crosslinks satisfied in the model of Nsp1 on the 40S subunit from being satisfied in the 43S-bound state (Fig. S4). Indeed, the contacts formed by Nsp1 on the surface of the 43S preinitiation complex clearly show that a single conformer cannot explain all observed crosslinks, and that multiple conformations of the 43S complex must also be present in solution. In particular, our crosslinks indicate multiple conformations within the eIF3 core relative to the 40S body, and strong dynamics for the eIF3I-eIF3B-eIF3G N-terminal region module relative to the mRNA entry cavity. Fluorescence-based measurements have shown that binding of Nsp1 to the small ribosomal subunit is modulated by initiation factors, with eIF3J competing with Nsp1 for binding to the 40S (Wang *et al*, 2021a).

Other direct Nsp1 interactions included eIF4B and the translational repressor PAIRB/SERBP1, although both of these PPIs were identified by a single crosslink. Interestingly, the dataset provided a glimpse on the extensive interactions formed by the highly dynamic PAIRB on the ribosome. As Nsp1 and PAIRB share a binding site in the mRNA entry channel, it is possible that PAIRB is displaced there by the Nsp1 CTD (Schubert *et al*, 2020), leading to a previously-uncharacterised “unplugged” PAIRB-bound conformation of the 40S subunit.

The crosslinks also identified the binding sites of some previously uncharacterized proteins, including the stress granule protein PRC2C, among the top enriched proteins from the AP-MS analysis, which is found crosslinked to eIF3A.

The extensive network of interactions formed by the Nsp1 NTD and linker regions provide insights into the targeting of initiation intermediates by Nsp1, and are consistent with the proposed role of these regions in host shutoff. These interactions may indeed provide a platform for the recruitment of the unknown downstream mRNA degradation machinery, for which we provide plausible candidates in our AP-MS study.

## Discussion

While the cryo-EM structures provided a clear mechanism for Nsp1-mediated host shutoff via its CTD, it has become clear that the Nsp1 NTD and linker are involved in additional mechanisms that allow for the escape of translational repression on viral transcripts and the targeting of host-translating ribosomes.

Our AP-MS results recapitulate the findings of cryo-EM, clarifying earlier results that did not detect an enrichment in ribosomes in the Nsp1-bound proteome, presumably because of the use of C-terminal tags, the high background of the ribosome, or the presence of nucleases in the preparation (Gordon *et al*, 2020; Stukalov *et al*, 2021). Proteomic analysis of ribosome populations further confirmed that full-length Nsp1 preferentially binds 40S and 43S complexes, which may be explained by a direct or RNA-mediated interaction between Nsp1 and eIF3G.

The crosslinking-MS data show that full-length Nsp1 is involved in a wide range of protein-protein interactions on the platform provided by the 40S subunit or the 43S preinitiation complex. We obtain structural information on these interactions via crosslinking-MS. Our results show that, despite its flexible position, the Nsp1 NTD interacts with the cavity facing the mRNA entry site on the 40S subunit and with the RRM domain of eIF3G. Moreover, the crosslinks further clarify current structural models of Nsp1-bound ribosomes and initiation complexes by unambiguously placing the eIF3G RRM and providing alternative conformations for the eIF3B-eIF3I module.

The interactions reported in our study provide context for recent observations that the NTD and linker regions are required for translational suppression. Indeed, K125, a residue required for 40S subunit binding in *in vitro* assays (Mendez *et al*, 2021), is found extensively crosslinked to eIF3G and several 40S subunits (Fig. 1A). In integrative models, this residue is located on the face of the Nsp1 NTD that interacts with RS3 (Fig. 1G). The model also accounts for the correlation of deletion of the 500-532 genome locus, located in the NTD β-sheet, with lower viral loads and non-severe infection traits (Lin *et al*, 2021). These residues may also be involved in contacts with the eIF3G RRM domain.

The dynamic nature of the Nsp1 linker is recapitulated in our data, as the large localization probability density for this region shows. Interestingly, NMR studies of Nsp1 in isolation (Wang *et al*, 2021b) have shown that the linker is partially but not completely disordered, which is consistent with our data showing a preference for a localization on the right-side of the ribosomal entry site cavity. Deletions in the linker residues K141-S142 in SARS-CoV Nsp1 impairs the translation shutoff function in CoV Nsp1 (Jauregui *et al*, 2013), indicating that this function may proceed downstream of ribosome binding. A similar deletion, K141-F143, has been observed in some SARS-CoV-2 patients (Benedetti *et al*, 2020). Our integrative model places these residues near RS3. The importance of the Nsp1 NTD and linker is underscored by their highly conserved nature between SARS-CoV species (Min *et al*, 2020; Thoms *et al*, 2020) and within SARS-CoV-2 lineages (Fig. S9).

Extensive modulation of eIF3G-RNA binding has been detected in surveys of RNA-protein in SARS-CoV-2 infected cells (Kamel *et al*, 2021; Schmidt *et al*, 2020), and directly implicated binding to viral factors. Upregulation of eIF3G and eIF4 RNA binding has been detected as a marker for the late-infection stage. Our crosslinking-MS network shows eIF3G crosslinked to the eIF4F subunits eIF4A, eIF4A2 and eIF4G1 of the eIF4F complex. No direct interaction between eIF4F and Nsp1 was observed.

In addition to its role in translational shutoff, Nsp1 is reported to block mRNA export from the nucleus via an interaction with the NXF1 export factor (Zhang *et al*, 2021). The required region for Nsp1-NXF1 interaction contains the NXF1 RRM domain, which may be further evidence of the ability of Nsp1 to interact with RRM domains.

Our model is consistent with results that indicate a competition between Nsp1 and eIF3J binding to the ribosome (Wang *et al*, 2021a). In eIF3J-bound, closed head conformations (Kratzat *et al*, 2021), the space occupied by the Nsp1 NTD in our main cluster of models clashes with the locked rRNA (Fig. S10). Indeed, eIF3J is less enriched than the core eIF3 complex in AP-MS, suggesting a preference for inhibiting open head conformations. The marked enrichment of the ABC-type ATPase ABCE1, found in some eIF3J-bound 43S states (Kratzat *et al*, 2021) is consistent with the presence of this factor in Nsp1-bound ribosomes (Thoms *et al*, 2020).

Our work explains the high conservation of the Nsp1 NTD and linker in coronaviruses. Far from being a simple “steric plug”, the NTD and linker are involved in an extensive network of protein-protein interactions. The model presented here also provides a framework for understanding the effects of previously reported deletions (Shi *et al*, 2020) in terms of protein conformational dynamics. We speculate that the interactions may partially explain the selectivity of Nsp1 for host-translating ribosomes, since translation initiation on a viral transcript may occur via an eIF3G-independent mechanism. Alternatively, the interaction to eIF3G may sequester this protein off of the ribosome. Nevertheless, it is clear that the Nsp1 NTD and linker provide an interaction platform for efficient translational repression and host shutoff.

## Materials and methods

### In-lysate crosslinking Mass spectrometry

#### Cell culture

HEK293T cells (DSMZ - ACC 635) were cultivated in Dulbecco’s modified Eagle’s Medium (DMEM, Corning), supplemented with 10% (volume/volume) fetal bovine serum (Sigma Aldrich) and 2 mM L-Glutamine (Corning) at 37°C and 5% CO_2_ in humidified atmosphere. Cells were seeded in 145 mm cell culture dishes and grown until 60-80% confluence. HEK293T cells were transfected with pcDNA5_FRT_TO-3xFLAG-3C-Nsp1 (Prof. Beckmann, Ludwig-Maximilians-Universität München). Transfection of HEK293T cells was performed by mixing plasmid DNA with PolyJet™ In Vitro DNA Transfection Reagent (SignaGen Laboratories) and incubation with cells for 3 h. Transfected cells were harvested after 48 hours.

#### In-lysate crosslinking and affinity purification

HEK293T cells were detached and washed twice with PBùS and harvested by centrifugation (400 g, 5 min). The cell pellet was lysed in lysis buffer [20 mM HEPES pH 7.5, 150 mM KOAc, 5 mM MgCl_2_, 1 mM DTT, 5% glycerol, 1x protease inhibitors and 0.5% NP-40] via sonication. Cell debris was pelleted by centrifugation. For in-lysate crosslinking a 50 mM stock of disuccinimidyl sulfoxide (DSSO, Cayman Chemical Company) was freshly prepared in Pierce™ Dimethylformamide (DMF, Thermo Scientific) and added to the lysate to a final concentration of 2 mM. The reaction was performed at RT for 1 h while gentile shaking and stopped using 1 M TRIS-HCl pH 7.5 to a final concentration 50 mM. ANTI-FLAG M2 agarose beads were washed with lysis buffer and added to the crosslinked lysate incubated for 1 h at RT while rotating. Beads were collected via centrifugation and washed twice with lysis buffer supplemented with 0.01% NP-40 and once with lysis buffer supplemented 0.05% NP-40. Washed beads were incubated with a buffer containing 20 mM HEPES-KOH pH 7.5, 150 mM KOAc, 5 mM MgCl_2_, 1 mM DTT, 0.05% NP-40. HRV 3C Protease (Thermo Scientific, Cat No. 88946) was added to the washed beads and incubated overnight at 4°C while rotating. Eluate was applied to crosslinked peptide preparation and enrichment.

#### Crosslinked peptide preparation and enrichment

Eluate from in-lysate crosslinking experiment was mixed with four times ice-cold Acetone (EMSURE®, Sigma Aldrich) and NaCl (to a final concentration of 150 mM) for protein precipitation. The mixture was stored at -20°C for 1 h and centrifuged and the supernatant was discarded. The dried protein pellet was solubilized in 8 M urea / 2 M thiourea solution, reduced using dithiothreitol (DTT) at 10 mM following incubation at RT for 30 min and derivatized at 30 mM iodoacetamide (IAA) over 20 min at RT and in the dark. Lys-C protease (MS Grade, Thermo Scientific) was added (in a ratio 1:100 (m/m) protease:protein) the samples were incubated for 2 h at 25 °C. Later the samples were diluted five times with 50 mM ammonium bicarbonate (ABC) and trypsin protease (MS Grade, Thermo Scientific) was added at a ratio of ∼1:50 (m/m). After 16 h at 25 °C the digestion was quenched by acidification with trifluoroacetic acid (TFA). Digested material was cleaned-up using C18 StageTips. Peptides were eluted and fractionated by size exclusion chromatography (SEC) using a Superdex Peptide 3.2/300 column (GE Healthcare, Chicago, IL, USA) at a flow rate of 10 μl min−1 using 30% (v/v) acetonitrile (ACN) and 0.1 % (v/v) TFA as mobile phase. Early 50-μl fractions were collected, dried and stored at −20 °C prior to LC-MS analysis. For linear identification of peptides in crosslinked samples 10% of each early fraction was pooled, dried and stored −20 °C prior to LC-MS analysis.

#### LC-MS protein identification of crosslinked samples and analysis

LC-MS/MS analysis of DSSO crosslinked sample was performed on an Orbitrap Fusion Lumos Tribrid mass spectrometer (Thermo Fisher Scientific, Germany) connected to an Ultimate 3000 RSLCnano system (Dionex, Thermo Fisher Scientific, Germany), which were operated under Tune 3.4, SII for Xcalibur 1.6 and Xcalibur 4.4. Fractions from peptide SEC were resuspended in 1.6% ACN 0.1% formic acid and loaded onto an EASY-Spray column of 50 cm length (Thermo Scientific) running at 300 nl/min. Gradient elution was performed using mobile phase A of 0.1% formic acid in water and mobile phase B of 80% acetonitrile, 0.1% formic. For each SEC fraction, we used an optimized elution gradient (from 2–18% mobile phase B to 37.5-46.5% over 90 min, followed by a linear increase to 45–55 and 95% over 2.5 min each). Each fraction was analyzed in duplicates. The settings of the mass spectrometer were as follows: Data-dependent mode with 2.5s-Top-speed setting; MS1 scan in the Orbitrap at 120,000 resolution over 400 to 1500 m/z with 250% normalized automatic gain control (AGC) target; MS2 scan trigger only on precursors with z = 3–7+, AGC target set to “standard”, maximum injection time set to “dynamic”; fragmentation by HCD employing a decision tree logic with (Giese *et al*, 2017) optimized collision energies; MS2 scan in the Orbitrap at a resolution of 60,000; dynamic exclusion was enabled upon a single observation for 60 s. Each LC-MS acquisition took 120 min.

#### LC-MS protein identification for generation of sequence database for crosslinking-MS

Protein identifications in pooled peptide SEC fractions from in-lysate crosslinking were conducted using a Orbitrap Fusion Lumos Tribrid mass spectrometer (Thermo Fisher Scientific, Germany) connected to an Ultimate 3000 RSLCnano system (Dionex, Thermo Fisher Scientific, Germany), which were operated under Tune 3.4, SII and Xcalibur 4.4. 0.1% (v/v) formic acid and 80% (v/v) acetonitrile, 0.1% (v/v) formic acid served as mobile phases A and B, respectively. For each experiment and replica 10 mAu from peptide SEC fractions 6-12 were collected dried and resuspended in 2% acetonitrile, 0.1% formic acid before injection onto an Easy-Spray column (C18, 50 cm, 75 μm ID, 2 μm particle size, 100 Å pore size) operated at 45 °C and running with 300 nl/min flow. Peptides were eluted with the following gradient: 0 to 2% buffer B in 2 min, 2 to 7.5% B in 5 min, 7.5 to 42.5%B in 80 min, 42.5 to 50% in 2.5 min. Then, the column was set to washing conditions within 3.5 min to 95% buffer B and flushed for another 5 min. For the mass spectrometer the following settings were used: MS1 resolution = 120,000; AGC target = 250%; maximum injection time = auto; scan range from 375 to 1500 m/z; RF lens = 30%. The selection criteria for ions to be fragmented were: intensity threshold > 5.0e3; charge states = 2–6; MIPS = peptide. Dynamic exclusion was enabled for 30 s after a single count and excluded isotopes. For MS2, the settings were: quadrupole isolation window = 0.4 m/z; minimum AGC target = 2.5 × 10e4; maximum injection time = 80 ms; HCD (higher-energy collisional dissociation) fragmentation = 30%. Fragment ion scans were recorded with the iontrap in ‘rapid’ mode with: mass range = normal. Each LC-MS acquisition took 120 min.

#### Generation of database for crosslink identification

Files from raw data obtained from linear identification of proteins were pre-processed using MaxQuant (1.6.17.) (Cox & Mann, 2008). Default settings with minor changes were used: three allowed missed cleavages; up to four variable modifications per peptide including oxidation on Met, acetylation on protein N-terminal. Carbamidomethylation on Cys was set as fixed. For the search ‘matching between runs’ feature was enabled with default settings. The full human proteome with 20.371 proteins plus amino acid sequence of Nsp1 was used. For protein quantification two or more peptides using the iBAQ approach were applied.

#### Crosslinking identification and validation

Raw files from crosslinking-MS acquisitions were converted to mgf-file format using MSconvert and were recalibrated to account for mass shifts during measurement. Recalibrated files were analyzed using xiSearch 1.7.5.1 (Mendes *et al*, 2019) with the following settings: MS1/MS2 error tolerances 2 and 4 ppm, allowing up to 2 missing isotope peaks (Lenz et al., 2018); tryptic digestion specificity with two missed cleavages; carbamidomethylation (Cys, +57.021464 Da) as fixed and oxidation (Met, +15.994915 Da) as variable modification, losses: –CH3SOH/ –H2O / –NH3, DSSO (158.0037648 Da linkage mass) with variable crosslinker modifications (“DSSO-NH2” 175.03031 Da, “DSSO-OH” 176.01433 Da). Additionally the maximum number of variable modifications per peptide was set to 1 and the additional loss masses were defined accounting for its cleavability (“A” 54.01056 Da, “S” 103.99320 Da, “T” 85.98264). Defined crosslink sites for DSSO were allowed for side chains of Lys, Tyr, Ser, Thr and the protein N-terminus. The database was composed of 400 Swiss-Prot annotated entries for *Homo sapiens* (Human) (taxon identifier 9606) with the highest abundance with the addition of the sequence of Nsp1. Results were filtered prior to FDR to matches having a minimum of three matched fragments per peptide, a delta score of > 10% of the match score and a peptide length of at least five amino acids. Additionally, spectral matches were prefiltered before FDR estimation to only those that had cleaved crosslinker peptide fragments for both peptides. Results were then to an estimated false-discovery rate (FDR) of 2% on residue-pair-level using xiFDR (version 2.1.5.2) (Mendes *et al*, 2019). The resulting estimated protein-protein interaction FDR was 6%. Self- and heteromeric-crosslinks were handled separately for FDR estimation. Crosslinking results were visualized in xiView (Graham *et al*, 2019).

### Affinity Purification Mass Spectrometry

#### Affinity purification of Nsp1 for AP-MS

HEK293T cells for the AP-MS were transfected with pcDNA5_FRT_TO-3xFLAG-3C-Nsp1 or pcDNA3 (control) in triplicates as described above. Each replica of HEK293T cells transfected with either pcDNA5_FRT_TO-3xFLAG-3C-Nsp1 or pcDNA3 (control) were trypsinized and collected by centrifugation and washed twice with phosphate-buffered saline (PBS, Corning). Cells were lysed in lysis buffer [20 mM HEPES pH 7.5, 150 mM KOAc, 5 mM MgCl_2_, 1 mM DTT, 5% glycerol, 1x protease inhibitors and 0.5% NP-40] via sonication (Bandelin Sonopuls). Cell debris was pelleted by centrifugation and cleared lysate was incubated with ANTI-FLAG M2 agarose beads (Sigma-Aldrich) for 120 min at 4°C while rotating. Later ANTI-FLAG M2 beads were washed twice with lysis buffer supplemented with 0.01% NP-40 and once with lysis buffer containing 0.05% NP-40. Pierce™ HRV 3C Protease (Thermo Scientific™) was used to cleave the Nsp1 from the ANTI-FLAG M2 agarose beads therefore the washed beads were incubated with a buffer containing 20 mM HEPES-KOH pH 7.5, 150 mM KOAc, 5 mM MgCl_2_, 0.05% NP-40 and 40 μg HRV 3C Protease at 4°C overnight.

#### Protein digestion and peptide size exclusion chromatography

Eluates from affinity purification were mixed with four times ice-cold Acetone (Sigma Aldrich) and NaCl for precipitation of proteins. The mixture was stored at -20°C for 1 h and later centrifuged the supernatant was removed. Dried protein pellet was solubilized in 8 M urea / 2 M thiourea solution, reduced using 10 mM DTT for 30 min at RT and derivatized with 30 mM IAA over 20 min at RT and in the dark. Digestion with Lys-C protease (Thermo Scientific) in a ratio 1:100 (w/w) was carried out for 2 h at 25 °C. The sample was diluted 5-fold in 50 mM ABC and trypsin protease was then added at a ratio of ∼1:50 (m/m). After 16 h at 25 °C the digestion was quenched by acidification with TFA. Peptides were cleaned up using C18 StageTips. Eluted peptides were fractionated using a Superdex Peptide 3.2/300 column (GE Healthcare) at a flow rate of 10 μl min−1 using 30% (v/v) ACN and 0.1 % (v/v) TFA as mobile phase. For each experiment and replica peptide-containing fractions were pooled and absorbance at 215 nm was quantified.

#### LC-MS protein identification of Nsp1 for affinity purification enrichment

AP-MS experiments were acquired using a Orbitrap Fusion Lumos Tribrid mass spectrometer (Thermo Fisher Scientific, Germany) connected to an Ultimate 3000 RSLCnano system (Dionex, Thermo Fisher Scientific, Germany), which were operated under Tune 3.4, SII and Xcalibur 4.4. 0.1% (v/v) formic acid and 80% (v/v) acetonitrile, 0.1% (v/v) formic acid served as mobile phases A and B, respectively. Samples were resuspended in 2% acetonitrile, 0.1% formic acid before injection onto an Easy-Spray column (C18, 50 cm, 75 μm ID, 2 μm particle size, 100 Å pore size) operated at 45 °C and running with 300 nl/min flow. Peptides were eluted with the following gradient: 0 to 2% buffer B in 2 min, 2 to 7.5% B in 5 min, 7.5 to 42.5%B in 80 min, 42.5 to 50% in 2.5 min. Then, the column was set to washing conditions within 3.5 min to 95% buffer B and flushed for another 5 min. For the mass spectrometer the following settings were used: MS1 resolution = 120,000; AGC target = 250%; maximum injection time = auto; scan range from 375 to 1500 m/z; RF lens = 30%. The selection criteria for ions to be fragmented were: intensity threshold > 5.0e3; charge states = 2–6; MIPS = peptide. Dynamic exclusion was enabled for 30 s after a single count and excluded isotopes. For MS2 the settings were: quadrupole isolation window = 0.4 m/z; minimum AGC target = 2.5 × 10e4; maximum injection time = 80 ms; HCD (higher-energy collisional dissociation) fragmentation = 30%. Fragment ion scans were recorded with the iontrap in ‘rapid’ mode with: mass range = normal. Each LC-MS acquisition took 120 min.

#### AP-MS analysis

Raw data from mass spectrometry were searched as described above using MaxQuant (1.6.17.). The obtained LFQ values for each replica were normalized according to the sum of absorbance units from peptide size exclusion chromatography. Enrichment analysis was performed in Perseus (1.5.6.0) (Tyanova *et al*, 2016). Normalized LFQ intensities were log-2 transformed and reproducibility was assessed (Fig. S and filtered to proteins detected in all three replicas in either +Nsp1 or control experiment (Figure S6). Missing values were then imputed based on a random selection from a normal distribution, downshifted by 1.8 standard deviations shrunk by a factor of 0.3. Significance was determined by performing a two-tailed t-test and imposing a significance cutoff threshold of FDR< 0.05 and ≥2-fold differential abundance on a log 2 scale. FDR was derived by permutation.

### Sucrose gradient analysis

#### Sucrose density gradient ultracentrifugation fractionation

HEK293T cells were transfected with pcDNA5_FRT_TO-3xFLAG-3C-Nsp1 or with pEGFP-C1 in triplicates as described above. Lysis and sucrose gradient ultracentrifugation as previously described (Kraushar *et al*, 2021). Briefly, cells were lysed in: 20 mM HEPES, 100 mM KCl, 10 mM MgCl_2_, pH 7.4, supplemented with 20 mM DTT, 0.04 mM Spermine, 0.5 mM Spermidine, 1x Protease Inhibitor cOmplete EDTA-free (Roche, 05056489001), 200 U/mL SUPERase-In RNase inhibitor (ThermoFisher, AM2694), 0.3% v/v IGEPAL CA-630 detergent (Sigma, I8896). Lysates clarified by centrifugation and the supernatants were applied to sucrose gradient ultracentrifugation. The ribosomal content was estimated by A_260_ optical density units. Gradients were linearized with a BioComp Gradient Master 107ip. Ribosomes were separated on 5 to 45 % sucrose density gradients prepared in Beckman Coulter Ultra-Clear Tubes 344057 (for LC/MS). Base buffer for sucrose solutions consisted of 20 mM HEPES, 100 mM KCl, 10 mM MgCl_2_, 20 mM DTT, 0.04 mM Spermine, 0.5 mM Spermidine, 1x Protease Inhibitor cOmplete EDTA-free (Roche), 20 U/ml SUPERase-In RNase inhibitor (ThermoFisher), pH 7.4, prepared with 5 & 45% sucrose (w/v). To linearized gradients with a BioComp Gradient Master 107ip. Equal optical density units from cleared cell lysates were loaded onto the gradient and centrifuged in a SW40 rotor (Beckman Coulter) for 5 hr, 4 C, 25000 rpm. Sucrose gradients were fractionated using a BioComp Piston Gradient Fractionator and Pharmacia LKB SuperFrac, with real-time A260 measurement by an LKB 22238 Uvicord SII UV detector recorded using an ADC16 PicoLogger and associated PicoLogger software. Fractions from one replica of Nsp1-overexpressing cells were precipitated using ice-cold 90% Ethanol (≥99,5 %, Carl Roth) and centrifuged. The supernatant was removed and the dried protein pellet was solubilized in 8 M urea / 2 M thiourea solution, reduced using DTT at 10 mM following incubation at RT for 30 min and derivatized at 30 mM IAA over 20 min at RT and in the dark. Digestion and StageTip cleanup were performed as for AP-MS samples.

#### LC-MS protein identification of sucrose fractions

Proteins from sucrose gradient fractions were acquired in a Q Exactive HF mass spectrometer (Thermo Fisher Scientific, Bremen, Germany) coupled to an Ultimate 3000 RSLC nano system (Dionex, Thermo Fisher Scientific, Sunnyvale, USA), operated under Tune 2.9, SII for Xcalibur 1.4 and Xcalibur 4.1. 0.1% (v/v) formic acid and 80% (v/v) ACN, 0.1% (v/v) formic acid served as mobile phases A and B, respectively. Samples were loaded in 1.6% acetonitrile, 0.1% formic acid on an Easy-Spray column (C18, 50 cm, 75 μm ID, 2 μm particle size, 100 Å pore size) operated at 45 °C and running with 300 nl/min flow. Peptides were eluted with the following gradient: 2 to 4% buffer B in 1 min, 4 to 6% B in 2 min, 6 to 37.5%B in 72 min, 37.5 to 42.5% in 5 min, 42.5% to 50% in 6 min, 50% to 90% in 3 min and hold at 90% for 7.5 min followed by 90 to 2%B in 23.5 min. For the mass spectrometer the following settings were used: MS1 scans resolution 120,000, AGC target 3 × 10^6^, maximum injection time 50 ms, scan range from 350 to 1600 m/z. The ten most abundant precursor ions with z = 2–6, passing the peptide match filter (“preferred”) were selected for HCD (higher-energy collisional dissociation) fragmentation employing stepped normalized collision energies (29 ± 2). The quadrupole isolation window was set to 1.6 m/z. Minimum AGC target was 2.50 × 10^4^, maximum injection time was 50 ms. Fragment ion scans were recorded with a resolution of 15,000, AGC target set to 1 × 105, scanning with a fixed first mass of 100 m/z. Dynamic exclusion was enabled for 30 s after a single count and included isotopes. Each LC-MS acquisition took 120 min. Raw data from LC-MS runs were searched as described above using MaxQuant (1.6.17.).

### Integrative modeling

#### Accessible interaction volume analysis

The 43S preinitiation complex structure (PDB ID 6zp4) (Thoms *et al*, 2020) was adjusted as follows: the RRM domain assigned to “protein X” in the deposited structure was assigned to the RRM domain of eIF3G, according to crosslinking-MS. The domain was built in MODELLER version 9.23 (Webb & Sali, 2016) and fitted in the density for “protein X” in the 43S preinitiation complex. The N-terminal stretch of eIF3G (residues 94-140), known to interact with the WD40 domain of eIF3I, was also built in MODELLER and placed in contact with eIF3I by homology to the structure of the yeast eIF3b-CTD/eIF3i/eIF3g-NTD subcomplex (PDB ID 4u1e) (Erzberger et al. 2014), and in accordance with crosslinking data (Fig. 3C).

Accessible interaction volume analysis was performed with DisVis (van Zundert & Bonvin, 2015) using the 40S complex bound to Nsp1 (PDB ID 6zlw) and the 43S preinitiation complex bound to Nsp1 (6zp4) as the fixed chain, and the structure of the Nsp1 NTD as the scanning chain, with a 1 Ågrid spacing and rotational sampling interval of 12.5 ° The allowed Cα-Cα distance for restraints was set between 2 and 30 Å.

#### Building blocks and representation

The structure of the human 40S complex bound to Nsp1 (PDB ID 6zlw) (Thoms *et al*, 2020) was used as the starting point for modeling the interactions of full-length Nsp1 to the 40S ribosome using crosslinking-MS restraints. Full-length Nsp1 was modeled using the structure of the N-terminal domain (PDB ID 7k3n) (Semper C., Watanabe N., Savchenko A., 2021) and the C-terminal domain in the Nsp1-bound 40S complex structure (PDB ID 6zlw). For integrative modeling, all protein regions not present in deposited structures, including the Nsp1 linker and unresolved ribosomal protein regions, were coarse grained as beads. Models were coarse grained as two rigid bodies: one comprising the 40S complex and the Nsp1 CTD (E148-G180) and another one comprising the Nsp1 NTD (M1-R124) and linker (K125-D147). The Nsp1 CTD position was derived from the structure of the Nsp1-bound 40S complex (PDB ID 6zlw) and kept fixed during sampling, while the rest of Nsp1 was left fully flexible. The rigid body boundaries, protein identifications, and coarse-graining levels are reported in table S4. The 8-residue N-terminal tag on Nsp1 was not included in the model. All regions not resolved in previous structures were kept fully flexible. **Sampling, scoring and model selection**. The building blocks described above (40S ribosome + Nsp1 CTD, full-length Nsp1) were used to derive an integrative model based on crosslinking-MS, known structures, homology models and physical restraints with the integrative modeling platform (IMP, version 2.15). The overall modeling workflow is described in fig. S1.

369 crosslinks (171 heteromeric) were used as distance restraints using the Bayesian CrossLinkingMassSpectrometryRestraint function in IMP.pmi with both psi (crosslink nuisance) and sigma (positional uncertainty) to be sampled. The inflection point in the scoring function was set to 21.0 Å Cα-Cα, and the weight was set to 1. The weight of connectivity restraints was set to 1 and that of excluded volume restraints was set to 2.

Sampling was performed by Replica Exchange Gibbs sampling in IMP 2.15, using 20 replicas in a temperature range between 1.0 and 10 in 8 independent runs with randomized initial configurations. A model was saved every 10 sampling steps, with each sampling step allowing for a 0.1 radian rotation and a 6 Å translation of each bead and rigid body. The number of frames was set to 15,000.

Scoring was performed as reported previously (Viswanath *et al*, 2017; O’Reilly *et al*, 2020) (Fig. S2). Briefly, a subset of 11,846 models of the 2,400,00 sampled configurations were selected based on score cutoffs (Crosslinking restraint < 336.0, Excluded volume restraint <22.5). Sampling exhaustiveness and completeness was assessed based on the criteria proposed in (Viswanath *et al*, 2017). Sampling precision was established at the point in which the root-mean squared deviation (r.m.s.d.) threshold at which representative models are no longer different from each other in a statistically significant manner (p>0.05) and with a small effect size (V<0.1) and more than 80% of models fall in clusters in the subsequent r.m.s.d.-based clustering procedures, yielding a sampling precision of 12.5 Å (Fig. S2C). The solutions were clustered based on Cα r.m.s.d. at a cutoff equal to the sampling precision. The precision in the main cluster of solutions, defined as the average r.m.s.d. to the cluster centroid model, was 5.4 Å. For all precision and clustering calculations, r.m.s.d. was computed on the Nsp1 NTD (residues 8-123) and linker not previously resolved in structures (residues 124-147). The centroid model in the main cluster is taken as the representative model. Localization probability densities were defined as the probability of any voxel (here, 5×5×5 Å^3^) being occupied by a specific region in model densities over the selected cluster, each of which is obtained by convolving superposed models with a Gaussian kernel (here, with a standard deviation of 20.0 Å).

## Acknowledgments

We thank Marchel Stuiver for valuable support with cell culture work and Nsp1 preparation. We also thank Prof. B. Glaunsinger and Dr. A. Mendez (Howard Hughes Medical institute, University of Berkley) for critical reading of the manuscript and Prof. R. Beckmann (Ludwig-Maximilians-Universität München) for kindly providing the Nsp1 construct.

## Author contributions

Conceptualization : AG,JR

AP-MS,crosslinking-MS, proteomics:FS

Sucrose gradients: MLK

Modeling/data analysis: AG

Supervision:CS,MLK,JR

Writing- initial draft: AG,FS,JR

Writing- editing and revision CS,MLK,JR

## Data availability

Mass spectrometry raw and processed data for both crosslinking-MS and proteomics experiments are deposited in JPost (Okuda *et* al, 2017) ProteomeXChange with identifiers XXX reviewer key XXX (AP-MS, sucrose gradient proteomics, linear IDs for crosslinking MS), XXX reviewer key: XXX (crosslinking-MS). Integrative model of Nsp1 on the 40S subunit is deposited in PDB-Dev with identifier ZZZ. Data and code for the integrative model is deposited in Zenodo with doi: XXX.

