## supplementary_information for "SARS-CoV-2 Nsp1 N-terminal and linker regions as a platform for host translational shutoff"

### Table S1: Crosslinks

### Table S2: Normalized AP-MS results

### Table S3: AP-MS enrichment analysis

### Table S4: Coarse graining and representations in integrative modeling.

Each region was-coarse grained at the resolutions described. The "rigid body" column indicates whether these proteins belong to the same or different building blocks (numbering is arbitrary). Regions without a pdb template are left fully flexible. A yellow highlight indicates a protein with crosslinks to Nsp1.

| Protein | UniProtID | Residues | Residues per Bead | Belonging to rigid body | PDB template |
|---|---|---|---|---|---|
| 18S RNA | - | all | 25 | 2 | 6zlw |
| Nsp1 | P0DTD1 | 1-15 | 5 | 1 | - |
| Nsp1 | P0DTD1 | 16-124 | 5 | 1 | 7k3n |
| Nsp1 | P0DTD1 | 125-147 | 5 | 1 | - |
| Nsp1 | P0DTD1 | 148-180 | 5 | 1 | 6zlw |
| RACK1 | P63244 | all | 25 | 2 | 6zlw |
| RL41 | P62945 | all | 25 | 2 | 6zlw |
| RS10 | P46783 | 1-97 | 25 | 2 | 6zlw |
| RS10 | P46783 | 98-165 | 25 | 2 | - |
| RS11 | P62280 | all | 25 | 2 | 6zlw |
| RS12 | P25398 | 1-9 | 25 | 2 | - |
| RS12 | P25398 | 10-132 | 25 | 2 | 6zlw |
| RS13 | P62277 | all | 25 | 2 | 6zlw |
| RS14 | P62263 | 1-16 | 25 | 2 | - |
| RS14 | P62263 | 17-151 | 25 | 2 | 6zlw |
| RS15 | P62841 | 1-14 | 25 | 2 | - |
| RS15 | P62841 | 15-134 | 25 | 2 | 6zlw |

| | | | | | |
|---|---|---|---|---|---|
| RS15 | P62841 | 135-145 | 25 | 2 | - |
| RS15a | P62244 | all | 25 | 2 | 6zlw |
| RS16 | P62249 | 1-7 | 25 | 2 | - |
| RS16 | P62249 | 8-146 | 25 | 2 | 6zlw |
| RS17 | P08708 | all | 25 | 2 | 6zlw |
| RS18 | P62269 | all | 25 | 2 | 6zlw |
| RS19 | P39019 | all | 25 | 2 | 6zlw |
| RS2 | P15880 | 1-55 | 25 | 2 | - |
| RS2 | P15880 | 56-276 | 25 | 2 | 6zlw |
| RS2 | P15880 | 277-293 | 25 | 2 | - |
| RS20 | P60866 | 1-16 | 25 | 2 | - |
| RS20 | P60866 | 17-119 | 25 | 2 | 6zlw |
| RS21 | P63220 | all | 25 | 2 | 6zlw |
| RS23 | P62266 | all | 25 | 2 | 6zlw |
| RS24 | P62847 | 1-125 | 25 | 2 | 6zlw |
| RS24 | P62847 | 126-133 | 25 | 2 | - |
| RS25 | P62851 | 1-41 | 25 | 2 | - |
| RS25 | P62851 | 42-113 | 25 | 2 | 6zlw |
| RS25 | P62851 | 114-125 | 25 | 2 | - |
| RS26 | P62854 | 1-102 | 25 | 2 | 6zlw |
| RS26 | P62854 | 103-115 | 25 | 2 | - |
| RS27 | P42677 | all | 25 | 2 | 6zlw |
| RS27a | P62979 | 1-77 | 25 | 2 | - |
| RS27a | P62979 | 78-156 | 25 | 2 | 6zlw |
| RS28 | P62857 | 1-7 | 25 | 2 | - |
| RS28 | P62857 | 8-68 | 25 | 2 | 6zlw |
| RS29 | P62273 | all | 25 | 2 | 6zlw |
| RS3 | P23396 | 1-227 | 25 | 2 | 6zlw |
| RS3 | P23396 | 228-243 | 25 | 2 | - |
| RS30 | P62861 | 1-47 | 25 | 2 | 6zlw |
| RS30 | P62861 | 48-59 | 25 | 2 | - |
| RS3A | P61247 | 1-20 | 25 | 2 | - |
| RS3A | P61247 | 21-233 | 25 | 2 | 6zlw |
| RS3A | P61247 | 234-264 | 25 | 2 | - |
| RS4X | P62701 | all | 25 | 2 | 6zlw |
| RS5 | P46782 | 1-15 | 25 | 2 | - |

| | | | | | |
|---|---|---|---|---|---|
| RS5 | P46782 | 16-204 | 25 | 2 | 6zlw |
| RS6 | P62753 | 1-230 | 25 | 2 | 6zlw |
| RS6 | P62753 | 231-249 | 25 | 2 | |
| RS7 | P62081 | 1-7 | 25 | 2 | - |
| RS7 | P62081 | 8-194 | 25 | 2 | 6zlw |
| RS8 | P62241 | all | 25 | 2 | 6zlw |
| RS9 | P46781 | 1-181 | 25 | 2 | 6zlw |
| RS9 | P46781 | 182-194 | 25 | 2 | - |
| RSSA | P08865 | 1-207 | 25 | 2 | - |
| RSSA | P08865 | 208-295 | 25 | 2 | 6zlw |

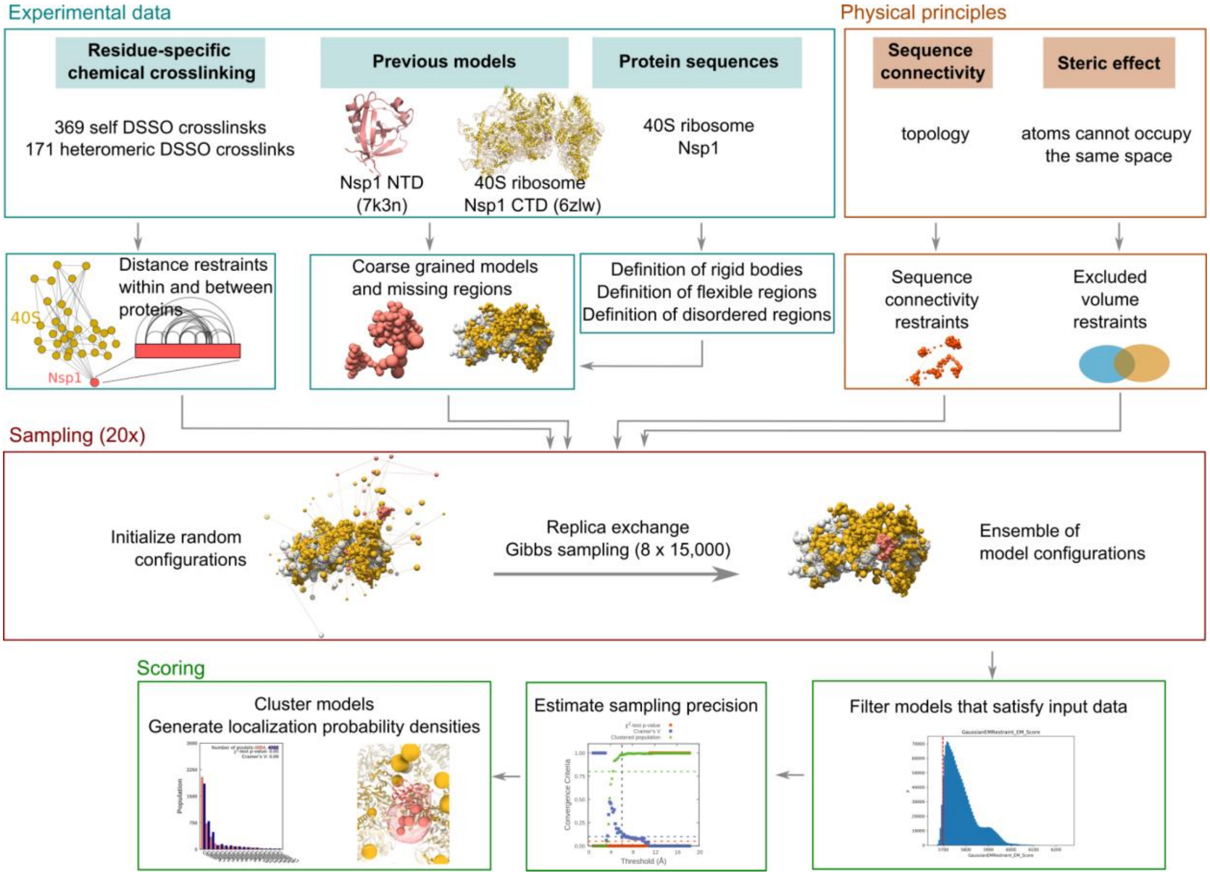


**Figure S1: Integrative modeling workflow.**

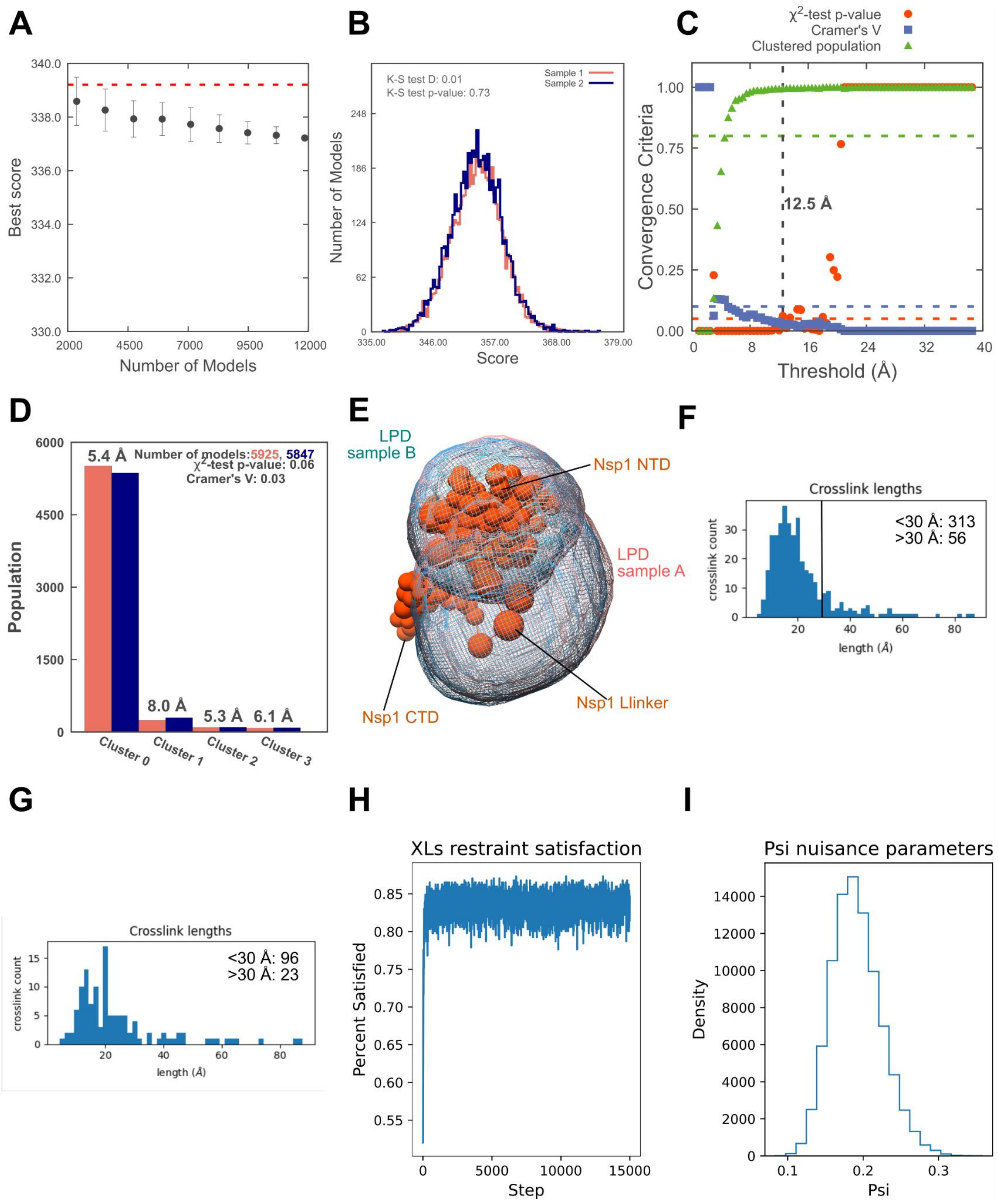


**Figure S2: Sampling completeness and precision of the integrative modeling procedure.**

**A)** Convergence of the model score in the 11,846 selected models. Minimum score of subsampling of random models is shown as mean +/- standard deviation (N=10). Score does not continue to improve as more models are sampled. **B)** Comparison of score distributions for the selected models in (A). The distributions (N=5956, 5890) display a negligible difference (D=0.01), and no statistically significant difference could be detected ($p>0.05$), indicating sampling completeness. Models in the first 4 modeling runs belong to sample 1, and models in the second 4 runs to sample 2. **C)** Estimation of sampling precision in the selected models as a function of r.m.s.d. sampling threshold. The vertical dashed line represents the threshold at which the p-value for homogeneity of proportions is satisfied ($p>0.05$ and the effect size is small (Cramer's V <0.1)) and >80% of models fall within one of the selected clusters. This indicates an overall sampling precision of 12.5 Å, as calculated on the Nsp1 NTD and linker regions. This is the smallest clustering threshold at which each sample contributes models proportionally to its size. **D)** Cluster populations obtained with an r.m.s.d. cutoff equal to the overall sampling precision. 91.8% of structures fall within one cluster, with a model precision (average r.m.s.d.-based distance to the cluster centroid model calculated over the ranges in Table S7) of 5.4 Å calculated on the Nsp1 NTD and linker regions. Colors refer to models originating from sample 1 or 2 in (B). **E)** Localisation probability densities for the Nsp1 NTD and linker in models from the two random samples that fall within cluster 0, indicating the in-cluster variance. Nsp1 in the cluster centroid model is shown in orange **F)** Crosslink distance distribution in the cluster centroid model. The vertical line marks 30 Å. The crosslink satisfaction is 85%. **G)** Crosslink distance distribution of crosslinks involving Nsp1 in the cluster centroid model. The vertical line marks 30 Å. The crosslink satisfaction is 80%, with violations consistent with the presence of a significant minor conformation in the proximity of RS2. **H)** Fraction of crosslinks having distance < 30 Å in a representative modeling run as a function of simulation frame. **I)** Density plot for crosslink uncertainty (Ψ), roughly equivalent to the sampled false discovery rate, during the 15,000 frames of a representative modeling run. The Ψ value peaks distribution indicates the presence of significant minor conformations, in agreement with visual inspection of the crosslinks.

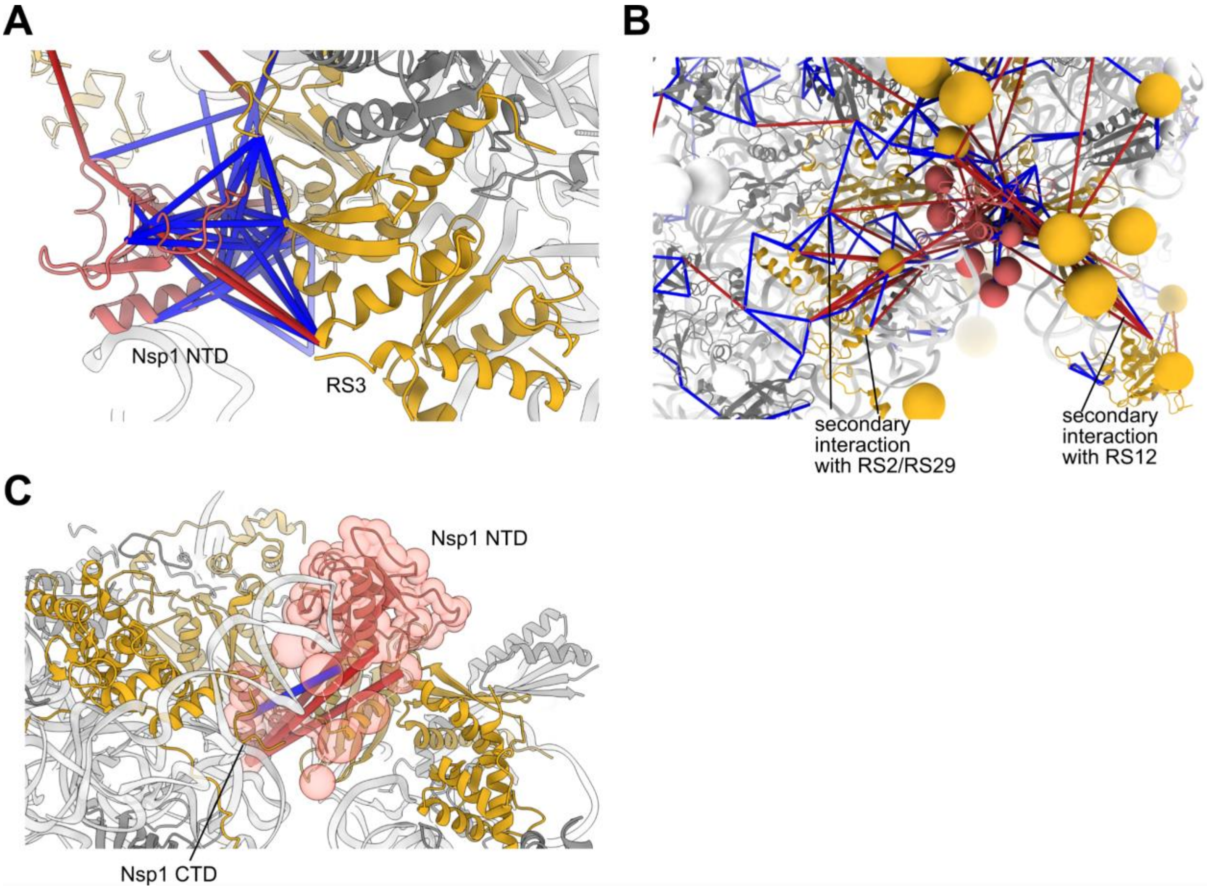


**Figure S3: Nsp1 conformation in main clusters of solutions.**

**A)** Interaction of Nsp1 with RS3 in the cluster centroid model. Satisfied crosslinks (<30Å) are shown in blue, violated crosslinks in red. Model precision is 5.4 Å. **B)** Position of the Nsp1 NTD in the cluster centroid model on the 40S subunit (pdb 6zlw). The consensus indicates a contact between Nsp1 and RS3, but crosslink violations are consistent with Nsp1 having secondary interactions on the other side and at the mouth of the mRNA entry channel cavity. The findings in solution reproduce the flexibility of the Nsp1 NTD and linker observed in cryo-EM structures. **C)** Self crosslinks between the Nsp1 CTD and NTD-linker regions. K164 faces away from RS2 and RS3 and is only observed crosslinked to the Nsp1 NTD and linker.

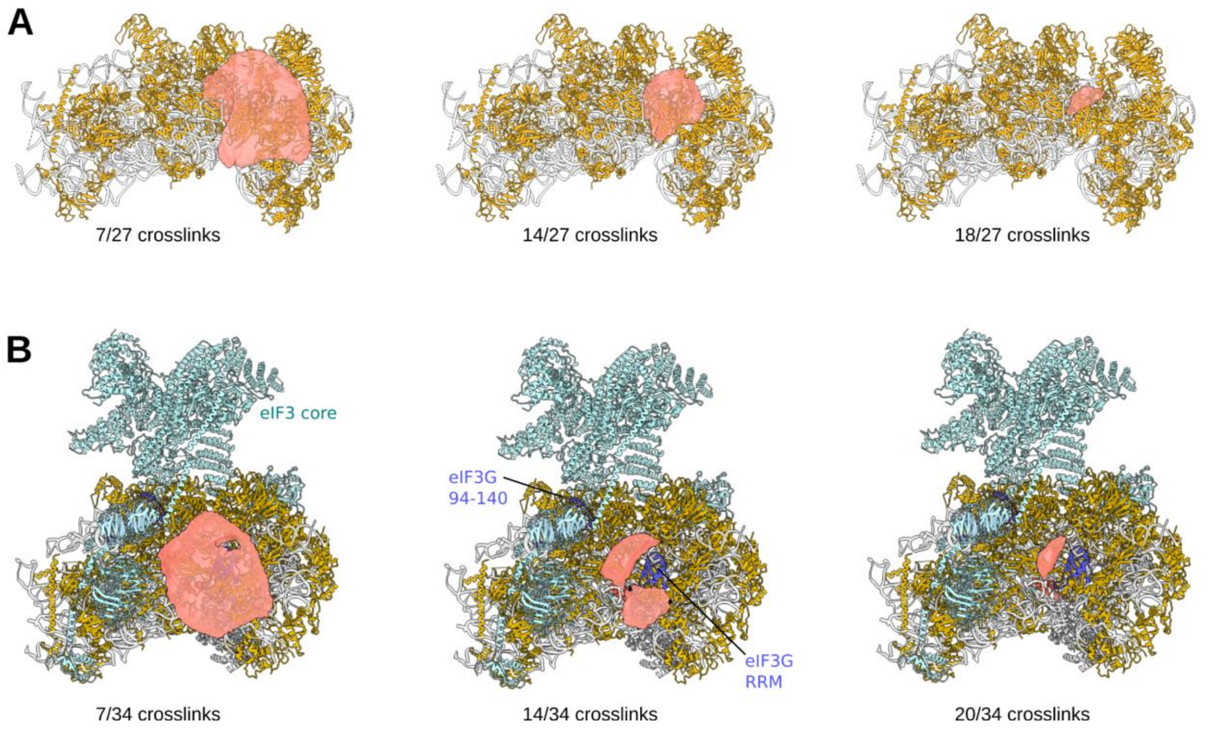


**Figure S4: Accessible interaction volume.**

**A)** Accessible interaction volume of the Nsp1 NTD on the 40S subunit (pdb 6zlw). The volume represents the positions of the center of mass of the domain consistent with at least 7, 14, and 18 crosslinks. The consensus indicates a contact between Nsp1 and RS3. Crosslinks to RS27A, RS9, RS17 and RS12 are inconsistent with the consensus, representing minor conformations, in accordance with the flexibility observed in cryo-EM. **B)** Accessible interaction volume of the Nsp1 NTD on the 43S complex captured in cryo-EM analysis (pdb 6zp4). The restraints between Nsp1 and eIF3G indicate that the NTD interacts with the eIF3G RRM domain. This interaction is however incompatible with direct binding to RS3 in this state.

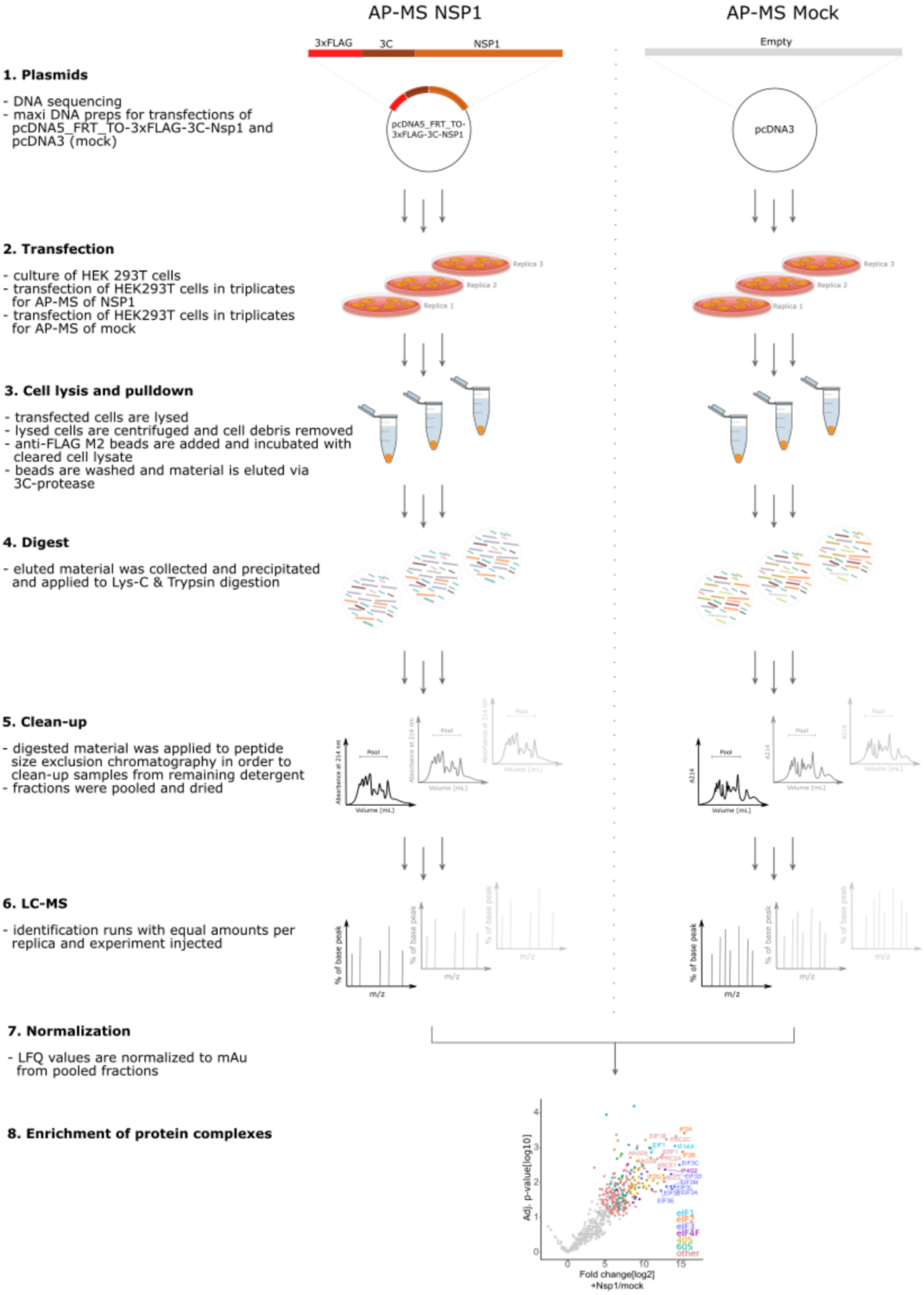
AP-MS NSP1
AP-MS Mock
3xFLAG
3C
NSP1
Empty
1. Plasmids
- DNA sequencing
- maxi DNA preps for transfections of pcDNA5_FRT_TO-3xFLAG-3C-Nsp1 and pcDNA3 (mock)
pcDNA5_FRT_TO-3xFLAG-3C-NSP1
pcDNA3
2. Transfection
- culture of HEK 293T cells
- transfection of HEK293T cells in triplicates for AP-MS of NSP1
- transfection of HEK293T cells in triplicates for AP-MS of mock
Replica 1
Replica 2
Replica 3
3. Cell lysis and pulldown
- transfected cells are lysed
- lysed cells are centrifuged and cell debris removed
- anti-FLAG M2 beads are added and incubated with cleared cell lysate
- beads are washed and material is eluted via 3C-protease
4. Digest
- eluted material was collected and precipitated and applied to Lys-C & Trypsin digestion
5. Clean-up
- digested material was applied to peptide size exclusion chromatography in order to clean-up samples from remaining detergent
- fractions were pooled and dried
Pool
Volume [mL]
6. LC-MS
- identification runs with equal amounts per replica and experiment injected
% of base peak
m/z
7. Normalization
- LFQ values are normalized to mAu from pooled fractions
8. Enrichment of protein complexes
Adj. p-value[log10]
Fold change[log2]
+Nsp1/mock
eIF1
eIF2
eIF3
eIF4F
40S
60S
other

**Figure S5: AP-MS workflow.**

Affinity purification of NSP1 for AP-MS. (1) Verification and DNA preparation of plasmids used for transfection of human cells. (2) Transfection of HEK293T cells in triplicates with FLAG-tagged NSP1 or with empty vector. (3) After 48h cells are harvested and lysed. For pulldown beads are added to cleared lysate and later 3C-protease is added to cleave bound material. (4) Eluted material was precipitated and applied to Lys-C and Trypsin digestion. (5) Peptide size exclusion chromatography was used to separate peptides from any contaminants originating from buffers used for pulldown. Early fractions were pooled and dried. (6) Equal amounts across replicates were used in identification runs. (7) LFQ intensities were normalized to the sum of absorbance units in peptide size exclusion chromatography. (8) Enrichment analysis of protein complexes.

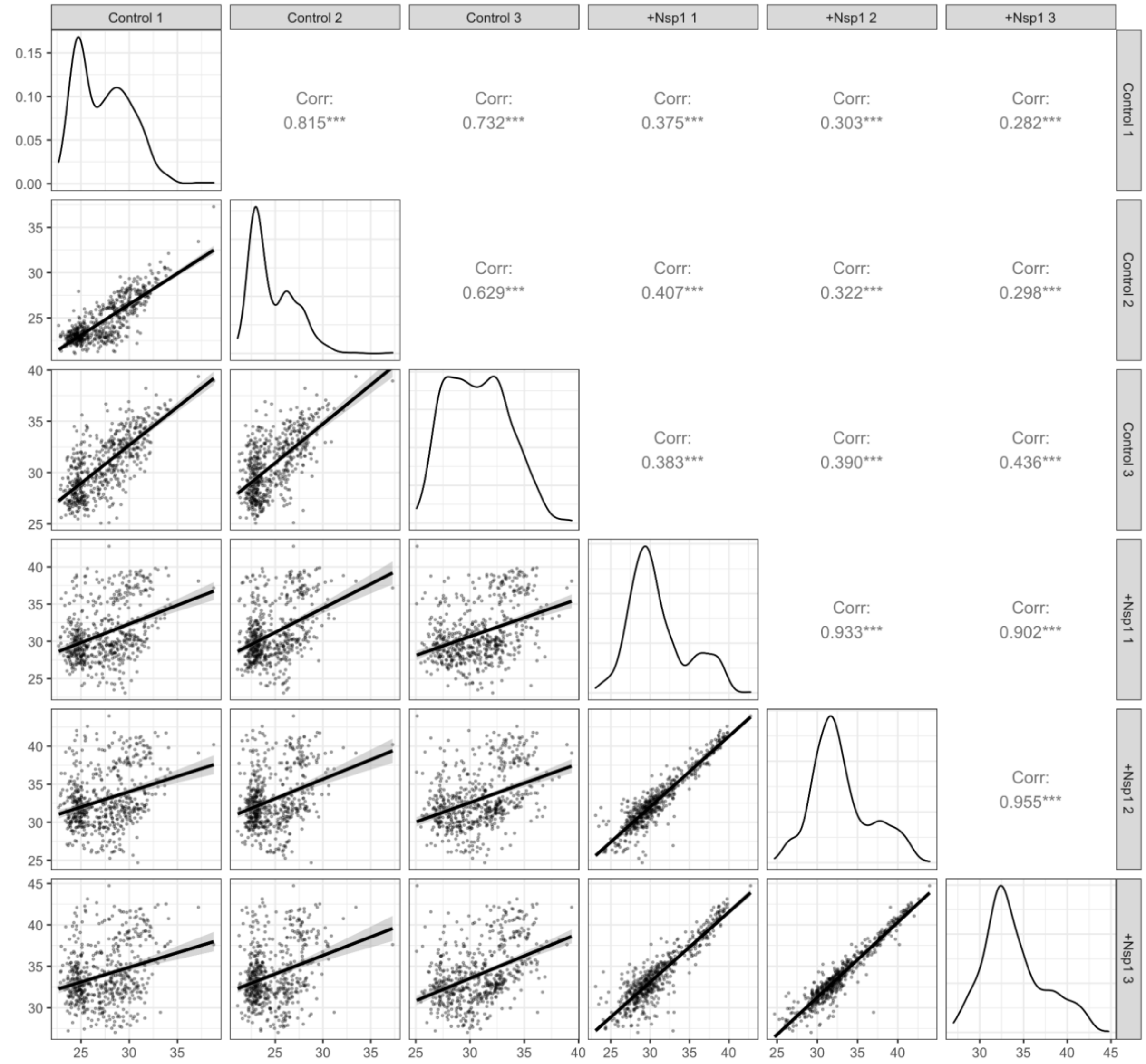


**Figure S6: AP-MS replicates.**

$Log_2$-transformed normalized LFQ intensities of AP-MS analyses of biological replicates of HEK293T cells transfected with pcDNA5_FRT_TO-3xFLAG-3C-Nsp1 (+Nsp1) or pcDNA3. Pearson correlation coefficient shown above diagonal with *** indicating p-value <0.001. Density plot along the diagonal. Scatterplot with linear regression and 95% confidence intervals shown below diagonal. Missing values imputed from normal distribution.

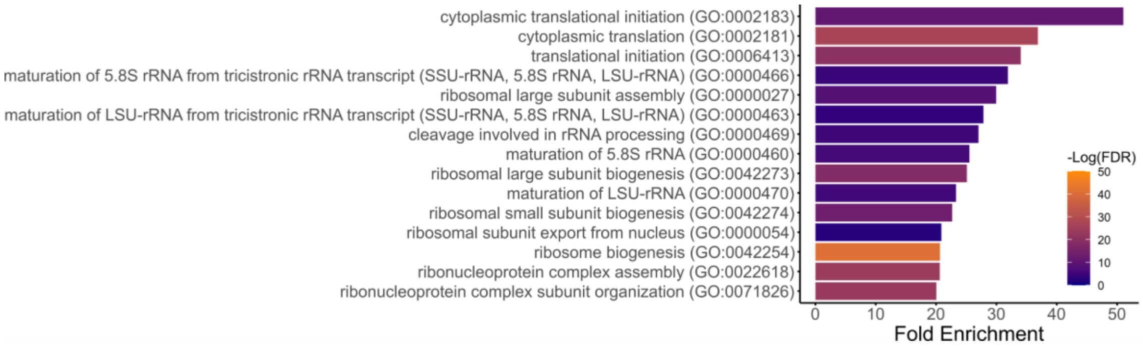


**Figure S7: Gene ontology enrichment analysis.**

GO-term enrichment analysis of proteome co-purifying with Nsp1, indicating translation initiation, overall translation and ribosome biogenesis among the most enriched biological processes. Fold enrichment and significance are measured relative to baseline proteome.

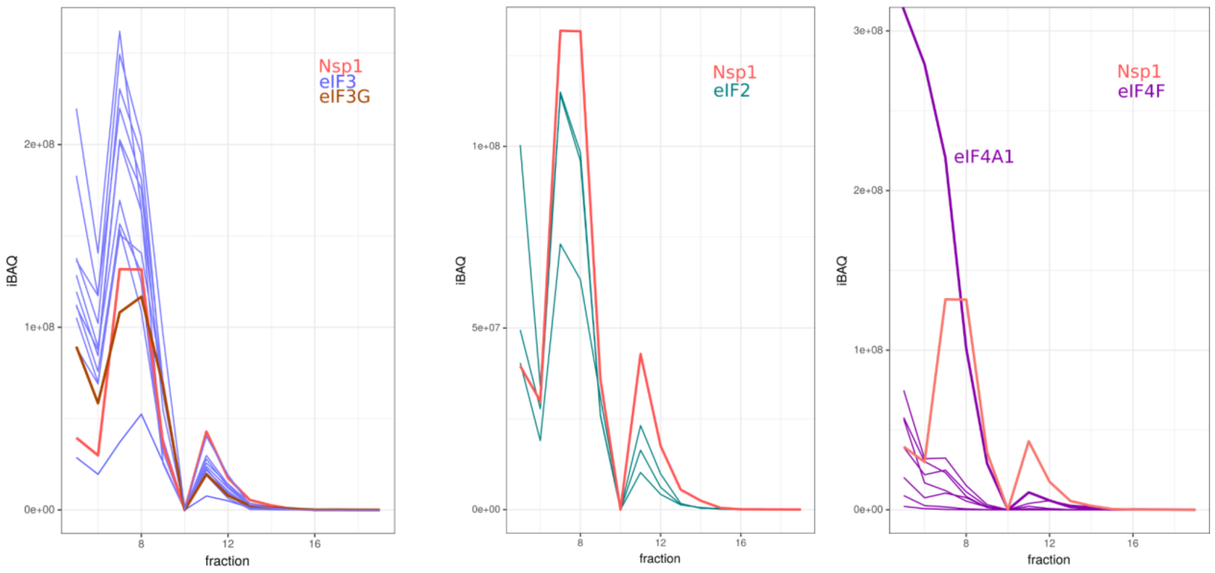


**Figure S8: Co-fractionation analysis of Nsp1 and eIF subunits.**

Proteomic analysis of sucrose gradient fractions from HEK293T cells expressing Nsp1. Nsp1 co-fractionates with components of the 43S complex, which elutes around fraction 8. A strong co-fractionation behavior is observed with eIF3G, which interacts with Nsp1 by crosslinking-MS. Weaker co-fractionation behavior is observed with the eIF4F complex.

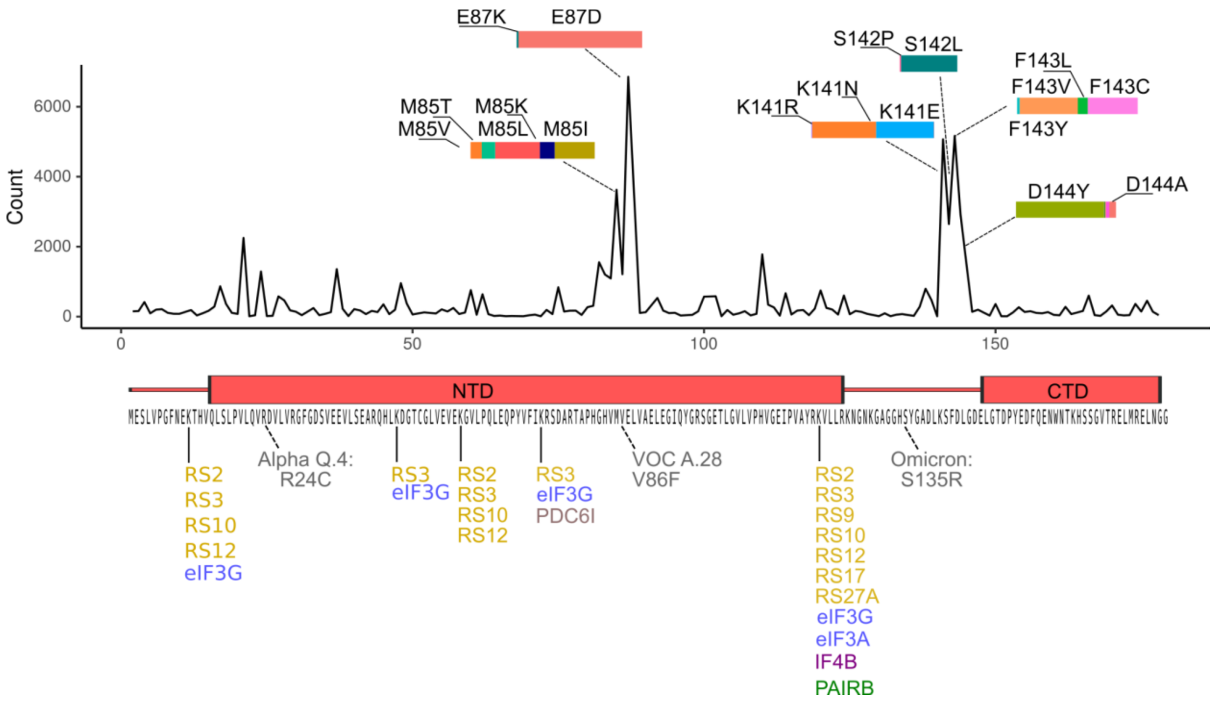


**Figure S9: Observed Nsp1 mutations.**

Top: Single nucleotide mutation counts for Nsp1 observed in the NCBI Sequence Read Archive. Data accessed via the NCBI Virus dashboard (https://www.ncbi.nlm.nih.gov/labs/virus/vssi/#/). SRA runs were selected which had at least 100 hits for SARS-CoV-2 via the SARS-CoV-2 Detection Tool, with a read length of at least 75, and were generated using the Illumina platform. Bottom: Nsp1 sequence and domain annotation. Solid lines indicate crosslinks observed in this study. Dashed lines indicate mutations observed in named variants, as described in the Stanford Coronavirus Antiviral & Resistance Database (https://covdb.stanford.edu/page/mutation-viewer/). Nsp1 is very stable, with low counts of non-synonymous mutations observed in the NCBI datasets and few mutations in reference SARS-CoV-2 lineages so far. Moreover, SARS-CoV-2 Nsp1 shares 84.4% sequence identity to SARS-CoV Nsp1.

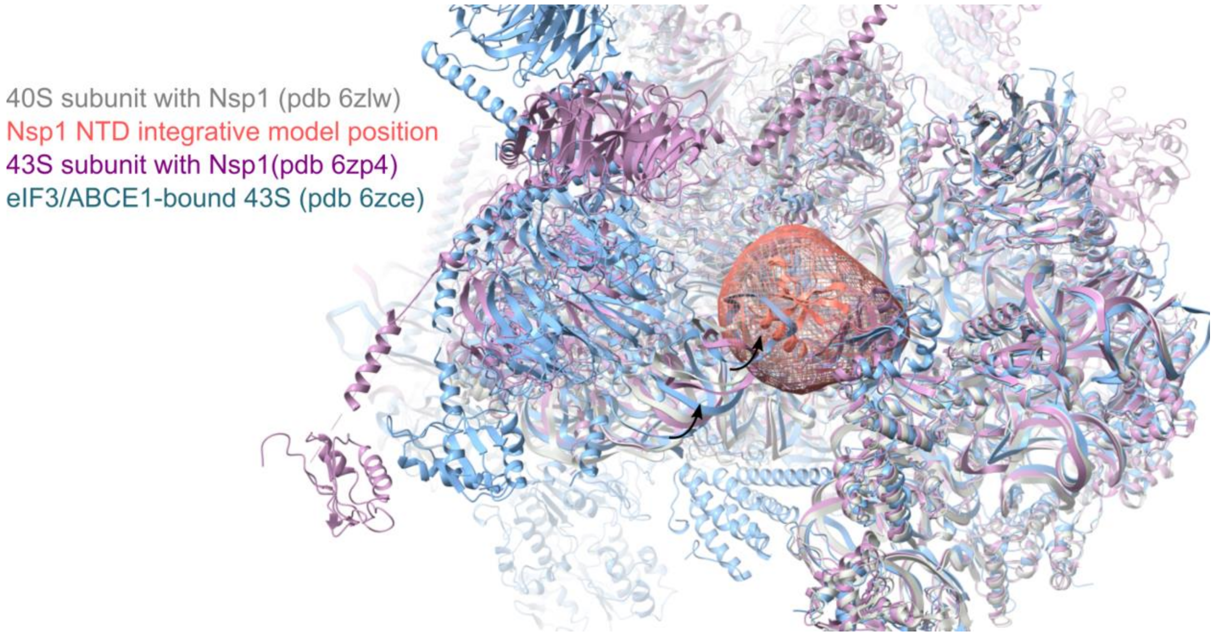


**Figure S10: eIF3J-bound 43S conformations clash with the position of Nsp1 on 40/43S subunit**

The main conformation of Nsp1 on the 40 and 43S subunit in the integrative is incompatible with the closed head conformation generated by eIF3J/ABCE1 binding due to steric clashes, consistent with competitive binding between Nsp1 and eIF3J to the 43S complex. The figure shows a superposition of the Nsp1-bound 40S, 43S structures with the Nsp1 in integrative model from this study and the yeast ABCE1/eIF3J-bound 43S preinitiation complex.
